# Genome-scale reconstruction of Gcn4/ATF4 networks driving a growth program

**DOI:** 10.1101/2020.01.29.924274

**Authors:** Rajalakshmi Srinivasan, Adhish S. Walvekar, Aswin Seshasayee, Sunil Laxman

## Abstract

Growth and starvation are considered opposite ends of a spectrum. To sustain growth, cells must manage biomolecule supply to balance constructive metabolism with high translation, through coordinated gene expression programs. Global growth programs couple increased ribosomal biogenesis with sufficient carbon metabolism, amino acid and nucleotide biosynthesis, and how this is collectively managed is a fundamental question. Conventionally, the role of the Gcn4/ATF4 transcription factor has been studied only in the context of amino acid starvation. However, high Gcn4/ATF4 has been observed in contexts of rapid cell proliferation, and the specific role of Gcn4 in growth contexts are unclear. Here, using a methionine-induced growth program in yeast, we show that Gcn4/ATF4 is the fulcrum through which metabolic supply dependent sustenance of translation outputs is maintained. Integrating time-matched transcriptome and ChIP-Seq analysis, we decipher genome-wide direct and indirect roles for Gcn4 in this growth program. Genes that enable metabolic precursor biosynthesis indispensably require Gcn4; contrastingly ribosomal genes are partly repressed by Gcn4. Gcn4 directly binds promoter-regions and transcribes a subset of metabolic genes, particularly driving lysine and arginine biosynthesis. Gcn4 also globally represses lys/arg enriched transcripts, which include the translation machinery. The sustained Gcn4 dependent lys/arg supply is required to maintain sufficient translation capacity, by allowing the synthesis of the translation machinery itself. Gcn4 thereby enables metabolic-precursor supply to bolster protein synthesis, and drive a growth program. Thus, we illustrate how growth and starvation outcomes are both controled using the same Gcn4 transcriptional outputs, in entirely distinct contexts.

## Introduction

Understanding the organizational principles of transcriptional programs that define growth or starvation is of fundamental importance. In order for cells to sustain growth, and thereby proliferation, a controlled supply of biosynthetic precursors is essential. These precursors include amino acids that drive protein translation, nucleotides (to make RNA and DNA), and several co-factors. Such a balanced cellular economy therefore requires coordinated, genome-wide responses in order to manage metabolic resources and ensure coordinated growth outputs. Here, the model eukaryote, *Saccharomyces cerevisiae*, has been instrumental in building our general understanding of global nutrient-dependent responses, addressing how cells allocate resources, defining transcriptional and metabolic ‘growth programs’, as well as to uncover general mechanisms of nutrient-sensing [1–9]. However, much remains unclear about how cells sustain the high requirement of biosynthetic precursors during growth programs.

Interestingly, studies from yeast and other systems show that the presence of some metabolites, even in nutrient-limited conditions, induces cell growth programs, as observed at the level of transcription, signaling or metabolism. One example is that of acetyl-CoA, which at sufficient concentrations induces cells to exit quiescence, and activates global gene expression programs driving proliferation [10–17]. Similarly, methionine (and its metabolite S-adenosyl methionine) turn on growth programs in cells [18–20]. In mammals, methionine availability correlates with tumor growth [21,22], and methionine restriction improves cancer therapy, by limiting one-carbon and nucleotide metabolism [23,24]. In yeast, supplementing methionine inhibits autophagy [25], activates growth master-regulators [18], and increases cell growth and proliferation [18,26]. At the level of global transcriptional and metabolic states, methionine triggers a hierarchically organized growth program, where cells transcriptionally induce ribosomal genes, and key metabolic nodes including the pentose phosphate pathway, as well as all amino acid, and nucleotide biosynthesis [20]. These are quintessential hallmarks of a cell growth program [27]. Therefore, using this controlled growth program, it may be possible to decipher universal regulatory features that determine a growth state. Further, such a system can be used to address how the metabolic program couples with the regulation of translation outputs. Unexpectedly, this previous study suggested that the transcription factor Gcn4 was critical for this growth program [20]. Such a role played by Gcn4 in a growth program was both unclear and unforeseen. This is because our current understanding of Gcn4 comes primarily from its role during starvation. Contrastingly, the role of Gcn4 during high cell growth is largely obscure.

Gcn4 (called ATF4 in mammals) is a transcriptional master-regulator, conventionally studied for its role during starvation and stress [28–31]. During severe amino acid starvation, the translation of Gcn4 mRNA increases, through the activation of the Gcn2 kinase, and subsequent eIF2-alpha phosphorylation [28,32,33]. This resultant increase in Gcn4 protein allows it to function as a transcriptional activator, where it induces transcripts involved in amino acid biosynthesis, thereby allowing cells to restore amino acid levels and survive starvation [28,31,34,35]. Almost our entire current knowledge of Gcn4 function comes from studying its roles during amino acid and other nutrient starvation. Contrastingly, we surprisingly found that in a growth program triggered by abundant methionine, cells induce Gcn4, in a context of high cell proliferation [20]. Other studies in several cancers suggest that the mammalian ortholog of Gcn4, called ATF4, is critical to sustain high growth [36,37]. Since starvation and growth programs are considered to be opposite ends of a spectrum, we wondered what specific roles does Gcn4 carry out during this growth program?

In this study, we find that Gcn4 controls essential components of an anabolic program, which are coupled with the management of overall translation. During such a growth program, Gcn4 directly transcribes genes required for amino acids and transamination reactions, and indirectly regulates essential ‘nitrogen’ metabolic processes, leading to nucleotide synthesis. We elucidate the direct and indirect, methionine-dependent roles of Gcn4, and identify separate requirements for this protein to control the metabolic component of this growth program, as well as manage the induction of translation-related genes. Thereby, we establish the importance of Gcn4-enabled biosynthetic precursor supply in appropriately maintaining a high translation capacity. Notably, comparing this function of Gcn4 during growth programs, to its well-known, conventional roles in starvation, reveals largely conserved transcriptional outputs of Gcn4 in both scenarios that however lead to distinct outcomes for the cell (growth vs survival). Through this, we show how a transcriptional master-regulator, conventionally viewed as a ‘survival-factor’, uses its canonical outputs to enable a growth program by ensuring specific amino acid synthesis in order to manage sufficient translation capacity.

## Results

### Methionine induces an universal ‘growth program’

Understanding the regulatory logic of transcriptional networks in growth programs is of fundamental importance. The role of the Gcn4 transcriptional master regulator has been well studied primarily in the context of severe nutrient starvation, as extensively explained in the subsequent section. However, several studies of cancers suggest that the mammalian ortholog of Gcn4 (ATF4) is required for rapid growth [36,37]. Therefore, we first wanted to establish a relevant, universal system where the role of Gcn4 during a growth program could be rigorously studied. Here, we utilized prior knowledge suggesting that methionine induces a transcriptional and metabolic growth program.

The observations showing that methionine switch cells to a growth state come primarily from yeast cells using lactate as a sole carbon source [18,20,25]. In these lactate-dependent conditions, global gene expression analysis revealed that providing methionine induces transcripts that represent a ‘growth signature’ [20]. This includes increased expression of ribosomal transcripts, and induced expression and metabolic flux through the pentose phosphate pathway, amino acid and nucleotide biosynthesis [20]. Since current studies are limited to only this lactate carbon source condition, we first more broadly established that this methionine response is universal, by studying the global transcriptional response to methionine supplementation in high glucose medium (the most preferred carbon source for yeast).

We performed comprehensive gene-expression analysis comparing transcripts from cells growing in glucose (MM) or glucose supplemented with methionine (MM+Met), as shown in Supplementary Figures 1 and 2, Supplementary WS2, Supplementary WS3, and described in the corresponding supplementary text. These results collectively show that the transcriptional response to methionine retains all the hallmarks of an anabolic growth program even when glucose is used as a carbon source. This includes the induction of appropriate metabolic genes (particularly all amino acid biosynthesis, nucleotide biosynthesis and transamination reaction related genes), as well as cytoplasmic translation related genes (Supplementary Figure 2, Supplementary WS2, Supplementary WS3). This transcriptional signature of cells MM+Met overlaps well with earlier studies of cells growing in lactate as a carbon source (supplemented with methionine) [20]. This induction of the translation machinery along with amino acid and nucleotide synthesis genes are all classic hallmark signatures of an anabolic program [27,38,39]. Further, these transcriptional changes also result in an appropriate metabolic state switch (increased *de novo* amino acid and nucleotide synthesis), as determined using a quantitative, targeted, stable-isotope pulsed LC/MS/MS based flux approach (Supplementary Figure 3).

In summary, we find that methionine triggers a growth program, with the induction of both metabolic and ribosomal genes, even in preferred medium with glucose as a carbon source. We therefore use this system (MM+Met) to address universal principles of cell growth regulation.

### GCN4 is induced by methionine and controls a conserved transcriptional signature in both growth and starvation programs

Conventionally, Gcn4 (a transcriptional master-regulator), is studied in the context of severe starvation, as part of the integrated stress response [28,29,31,40,41]. Nearly all existing studies of Gcn4 use pharmacological inhibitors of amino acid biosynthesis, such as 3-amino triazole (3-AT) or sulfo meturon (SM) to induce Gcn4, and study its role in starvation responses where cell growth is minimal [31,34,35,42,43]. Indeed, our current understanding of Gcn4 function comes primarily from contexts of nutrient-stress and starvation. In contrast, we had earlier observed that supplementing methionine strongly induces Gcn4 [20], coincident with *increased* cell growth and proliferation. Since this is distinct from conditions of starvation and low growth, we wanted to understand what the role of Gcn4 was, during a growth program.

We therefore used the methionine-induced growth transcriptional program (as described in Supplementary Figure 2 and the previous section) to address this question. We first asked if Gcn4 protein is induced in methionine-supplemented glucose medium. Indeed, Gcn4 protein levels substantially increase when methionine is supplemented (MM+Met) (Figure 1A and Supplementary Figure 4A). This observation reiterates that Gcn4 can be induced by growth signals (methionine) irrespective of carbon source. We therefore dissected how much of this anabolic program is mediated by Gcn4. To address this, we compared transcriptomes of wild type and *Δgcn4* cells in MM+Met (Supplementary Figure 4B), and found a striking, Gcn4-dependent global response in the presence of methionine. ∼900 genes were differentially expressed in Δ*gcn4*, compared to wild type cells in MM+Met. Here, 514 genes were upregulated, and 398 genes were downregulated in *Δgcn4* cells in the presence of methionine (fold change cut-off of >= 2 fold) (Supplementary Figure 4B & Supplementary WS1). As a control, in only MM medium (without supplemented methionine), far fewer genes (∼160) showed any differential expression at all in *Δgcn4* relative to WT (Supplementary Figure 4B). These data show that Gcn4 has a critical role for the methionine-dependent growth program in glucose.

**Figure 1:**
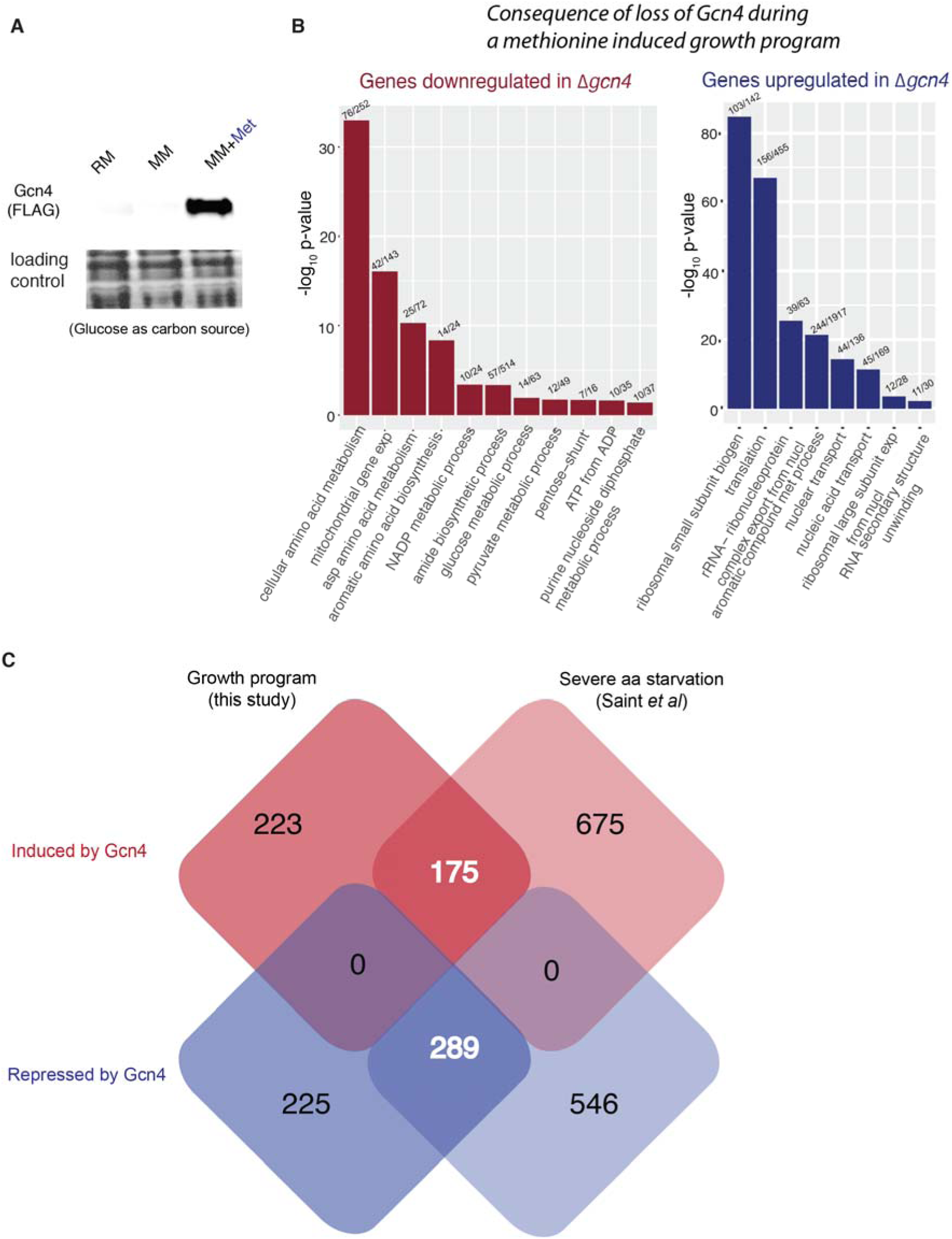
Gcn4 binds to its target gene promoters related to metabolism during a growth program. A. Gcn4 is strongly induced by methionine addition. A representative western blot shows high Gcn4 protein levels in MM+Met (Gcn4 tagged with the FLAG epitope at the endogenous locus). RM - rich medium, MM - minimal medium without amino acids and with glucose as a carbon source, and MM+Met - minimal medium without amino acids and with glucose as a carbon source supplemented with 2mM methionine. Also see Supplementary Figure 4A. B. GO based analysis and grouping of transcripts down regulated in *Δgcn4.* All the terms shown here are significantly enriched terms with the corrected p-value < 0.05 (hypergeometric test, Bonferroni Correction). GO based analysis and grouping of the transcripts up-regulated in *Δgcn4.* All the terms shown here are significantly enriched terms with the corrected p-value < 0.05 (hypergeometric test, Bonferroni Correction). Also see Supplementary WS3 and Supplementary Figure 4B for gene expression volcano plots. C. The Venn diagram shows the number of differentially expressed genes that overlap, from data obtained from distinct cell growth conditions where Gcn4 levels are high. The boxes on the left are data from this study (methionine induced growth program), while the boxes on the right use data from a severe amino acid starvation condition (sulfo meturon addition) [46]. Also see Supplementary WS6.

To understand the global consequences of the loss of Gcn4 during this growth program, we used a GO-based analysis to categorize most altered groups of genes. Upregulated genes in *Δgcn4* show a notable enrichment for ‘cytoplasmic translation’, ‘ncRNA processing’, ‘RNA maturation’ and ‘RNA methylation’ (Figure 1B). This strikingly revealed that the transcripts associated with protein translation, which were already induced by methionine, further increase in the absence of Gcn4. i.e. Gcn4 partially represses cytoplasmic translation even in a growth program. In contrast, genes that are downregulated in *Δgcn4* cells are primarily involved in amino acid biosynthetic processes, nucleotide biosynthetic processes, mitochondrial translation, NADP metabolic processes and pyruvate metabolism (Figure 1B & Supplementary WS3). Collectively, this reveals that Gcn4 is essential for the induction of genes involved in these metabolic processes, which is a majority of the methionine induced anabolic program, but partially represses translation.

Here, we note a striking observation. In studies of starvation, the induction of Gcn4 represses ribosomal genes, and induces amino acid biosynthesis genes [35,44,45]. In contrast, in this methionine-induced growth program, ribosomal and translation related genes are themselves induced even as Gcn4 is also induced. The loss of Gcn4 further increases ribosomal genes, suggesting that Gcn4 appropriately keeps the extent of ribosomal gene induction in check, while the ribosomal gene induction occurs through independent regulation. Furthermore, the induction of amino acid biosynthetic genes remains regardless of starvation or growth programs. This hints that despite growth and starvation being at opposite ends of a spectrum, the role of Gcn4 in either state might be conserved. To further address this possibly conserved global role of Gcn4, we compared the overlap of Gcn4 dependent, induced or repressed genes in this growth program, with existing data from a conventional starvation program where Gcn4 has high activity. This gene expression data comes from a conventional mode of inducing Gcn4, via inhibiting amino acid biosynthesis using a chemical inhibitor of amino acid biosynthesis (sulfometuron or SM) [46]. Notably, we find that 44% of the genes activated by Gcn4 and 56% of the genes repressed by Gcn4 in the methionine dependent growth program overlap with the genes activated and repressed in the SM dependent starvation condition (Fisher exact test, p<10^−10^) (Figure 1C). A GO grouping of the genes which overlap between the growth and the starvation condition suggests a conserved role of Gcn4 in inducing amino acid biosynthetic genes and in repressing translation related genes (Supplementary WS6).

Two key points emerge from these analyses. First, the role of Gcn4 appears to be conserved regardless of whether cells are in a growth or starvation program. This conserved role appears to be to increase transcripts related to amino acid and nucleotide biosynthesis (all required for anabolism), while repressing translation related genes. However, during a growth program, there is already an induction of translation genes (as seen in Figure 1). Therefore, in this context, Gcn4 tempers the extent of induction of translation related genes, while during starvation Gcn4 represses ribosomal genes below that of non-starved cells.

### Gcn4 binds to its target gene promoters related to metabolism during a growth program

Which parts of the transcriptional outputs in this growth program does Gcn4 directly regulate, and how does this compare to the known, direct roles of Gcn4 during starvation? To address this, we performed chromatin immunoprecipitation (ChIP)-sequencing of Gcn4 in MM and MM+Met conditions. Notably, this uniquely integrates directly comparable information from the Gcn4 ChIP-seq, with a time-matched global transcriptome, *during a growth program*.

First, we asked what is the Gcn4 DNA binding activity when induced by methionine. We performed ChIP of Gcn4 (with a FLAG-epitope incorporated into the C-terminus in the endogenous *GCN4* locus), using cells grown in MM and MM+Met, with MM essentially acting as a control. We considered peaks that are represented in both the biological replicates for further analysis, using very well correlated biological replicates (Supplementary Figure 4C). Here, we identified 320 Gcn4 binding peaks in the cells grown in MM+Met, whereas, there were no consensus peaks observed in replicate samples of cells grown in MM (Figure 2A & Supplementary WS4). The enhanced Gcn4 occupancy on the target gene promoter in MM+Met condition was further validated using ChIP-qPCR analysis (Supplementary Figure 6). This shows that the GCN4 occupancy on DNA increases in the presence of methionine.

**Figure 2:**
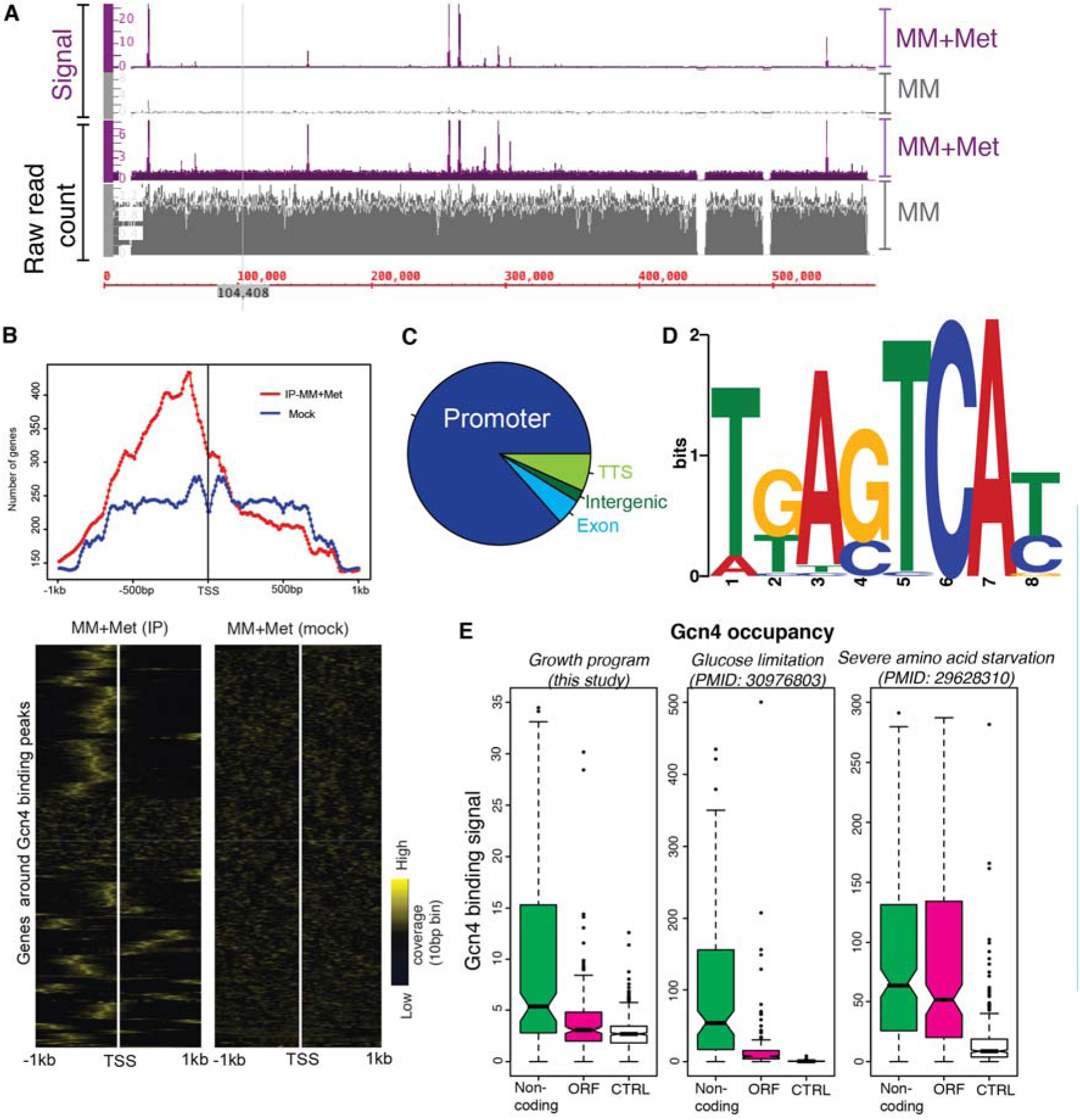
Gcn4 binds to its target gene promoters related to metabolism during a growth program. A. Genomic tracts showing Gcn4 binding to DNA regions in MM and MM+Met. Raw read counts and signal around the binding region of Gcn4 are shown. B. (Top) Density plots showing that most target genes have Gcn4 binding peaks upstream of the Transcription Start Site (TSS) in the ChIP samples (red), whereas no such enrichment of genes is observed in mock samples (blue). (Bottom) A heat map showing read coverage for Gcn4 binding, including 1kb upstream and downstream of predicted/known transcription start sites (TSS) of target genes. All the genes that fall in the vicinity of 750bp around the identified Gcn4 binding peaks are considered to be target genes. The heat map on the left shows read coverage in IP samples, and on the right shows coverage in mock-IP (control) under in MM+Met condition. Also see Supplementary Figure 5 which shows read coverage for Gcn4, in the context of the translation start site (ATG) of each gene. C. A pie chart showing the genomic features of the identified peaks annotated using ‘annotatepeak’ function of Homer tool [48]. D. Consensus binding motifs identified in Gcn4 binding peaks from MM+Met conditions. E. Boxplots, showing the Gcn4 binding signal corresponding to different genomic features, under distinct growth scenarios. For Gcn4 binding in non-coding and open reading frame (ORF) regions (as reported in a previous study) [34], we compared the Gcn4 binding signal in the Gcn4 ChIP sequencing data from cells in MM+Met (current study), or under different starvation conditions (severe AA-starvation [34], or in glucose limitation [52]. Notably, in either MM+Met or during glucose limitation the Gcn4 binding signal in ORF peaks is significantly lower than the non-coding region peaks (p<10^−8^). Contrarily, under severe amino acid starvation [34], the Gcn4 binding signal found in ORF and non-coding regions are very similar.

Next, we analyzed the Gcn4 binding signals around the transcription and translation start site of the genes found within 750bp around the identified peaks. Transcription start site data available for cells growing in rich, glucose medium (the nearest possible condition to that used in this study) was obtained from the YeasTSS database [47]. The TSS identified using the CAGE method reported in this database was used for our analysis. Notably, a majority of the Gcn4 binding peaks in MM+Met are found upstream of these annotated transcription start sites (Figure 2B). A similar analysis with the translation start sites of the target genes shows higher read coverage upstream of the translation start site (Supplementary Figure 5). We further analyzed the genomic features of the identified peaks using the HOMER program [48]. Notably, we observed a very apparent enrichment of Gcn4 binding to the promoter region of the targets. 263 out of 320 peaks are found within the promoter region of target genes (−1kb to +100bp around the TSS), while the remaining peaks bind at intergenic regions (11), exons (17) or close to transcription termination sites (29) (Figure 2C & Supplementary WS4). This shows that during a growth program, Gcn4 activity is primarily restricted to binding promoter sites of target genes.

We next searched for the enrichment of sequence motifs in the peaks identified in the MM+Met condition using The MEME-suite [49]. We found that these peaks were enriched for the conserved Gcn4 binding motifs found previously under amino acid starvation conditions, [35,50,51]. Strikingly, 81% (260 out of 320) of the peaks that we identify have at least one of the variants of the Gcn4 binding motif ‘TGANTCA’ (Figure 2D), showing that Gcn4 in this context still primarily recognizes its high-affinity DNA binding motif.

Finally, how does this compare to studies of Gcn4 activity during starvation, particularly during severe amino acid biosynthesis inhibition [34,35]? A comprehensive previous study of Gcn4 function during amino acid starvation indicated substantial Gcn4 binding to regions within ORFs of genes, as well as to promoter regions [34]. To compare this study from a starvation program with our data from a growth program, for the non-coding and the ORF peaks regions reported in the previous study [34], we calculated the Gcn4 binding signal in our Gcn4 ChIP seq data (from the MM+Met condition). Strikingly, we find that the signal in ORF peaks is significantly lower than the non-coding peak under MM+Met condition (p-value < 10^−8^), whereas a similar analysis performed using the Gcn4 ChIP-seq data from [34] show little differences in the signal intensity between ORF and Non-coding peaks (p-value of 0.002) (Figure 2E). As a distinct comparison, we used a dataset from a milder starvation regime [52], where glucose was limited in a chemostat. Here, the occupancy of Gcn4 was more similar to that observed during our growth program, with a majority of Gcn4 occupancy at promoter regions of target genes (Figure 2E). These analyses show that during a growth program, the direct targets of Gcn4 remain highly specific, conserved and restricted to the promoter regions of genes. The Gcn4 occupancy limited to promoter regions during growth and mild starvation conditions can be possibly explained by a lower dosage of Gcn4 under these conditions. Under extreme amino acid starvation conditions, very high Gcn4 levels might result in increased Gcn4 occupancy on the ORFs, in addition to its specific binding to the promoter. Collectively, our data shows that regardless of the mode of Gcn4 induction, and whether cells are in a growth or starvation program, it binds specifically to a highly conserved motif.

Thus, the global role of Gcn4 during either a growth program, or in a starvation response appears remarkably conserved. However, the cellular outcomes are different, and this can be explained by two criteria. First, the amounts of Gcn4 protein (as induced by methionine) will be different from the other conditions tested, as the mode of induction of Gcn4 is entirely different in these studies. Therefore, since any protein’s affinity to its target depends on its dosage in the cell as well as the presence of other competing factors, there will be differential binding affinity to the targets, as is well known for most transcription factors [53,54]. Second, the context of Gcn4 induction is entirely distinct. In this context Gcn4 is supporting an anabolic program, while the cells also have increased ribosomal genes. Hence, while the function of Gcn4 is the same (primarily to induce amino acid biosynthesis, and indirectly repress translation), the outcome is entirely different, because in a growth program the increased production of amino acids and nucleotides might aid the increase in translational capacity via increased ribosomal biogenesis.

### Direct and Indirect targets of Gcn4 during a growth program

We therefore asked how much of the Gcn4-dependent transcriptional response is directly regulated by Gcn4, and what its specific targets were? To identify direct targets of Gcn4 in methionine-dependent gene regulation, we overlaid the transcriptome data (*Δgcn4 vs WT* in MM+Met) with the ChIP-seq data (from MM+Met). Out of the 398 genes that are downregulated in Δ*gcn4*, 133 are direct targets of Gcn4 (Supplementary WS4). Contrastingly, Gcn4 directly regulates only 24 out of 514 upregulated genes (Supplementary Figures S7A and S7B, and Supplementary WS4). These results strengthen the role of Gcn4 as a transcriptional activator. GO-based analysis of the genes directly transcribed by Gcn4 reveals a significant enrichment of amino acid biosynthetic genes. Notably, the indirectly activated targets are enriched for nucleotide biosynthesis, the pentose phosphate pathway, and mitochondrial translation (Figure 3A, Supplementary WS3). In addition to the amino acid biosynthetic genes, Gcn4 directly activates genes involved in other critical functions, particularly the Sno1 and Snz1 genes (pyridoxal synthase), required for transamination reactions that lead to amino acid synthesis, and Nde1-the NADH dehydrogenase (Supplementary Figure 7C). These genes have Gcn4 binding sites in its promoter [55]. In contrast to the Snz1 and Sno1 pair that is bidirectionally activated by Gcn4, the Trm1 and Mdh2 pair of genes are bidirectionally repressed by Gcn4 (Supplementary Figure 7D). These data show that in the presence of methionine, Gcn4 directly increases the expression of primarily the amino acid biosynthetic arm, whereas the methionine-dependent activation of nucleotide biosynthetic genes, pentose phosphate pathway, mitochondrial translation related genes are indirectly regulated by Gcn4. Collectively, the metabolic component of the methionine-dependent growth program is directly regulated by Gcn4.

**Figure 3:**
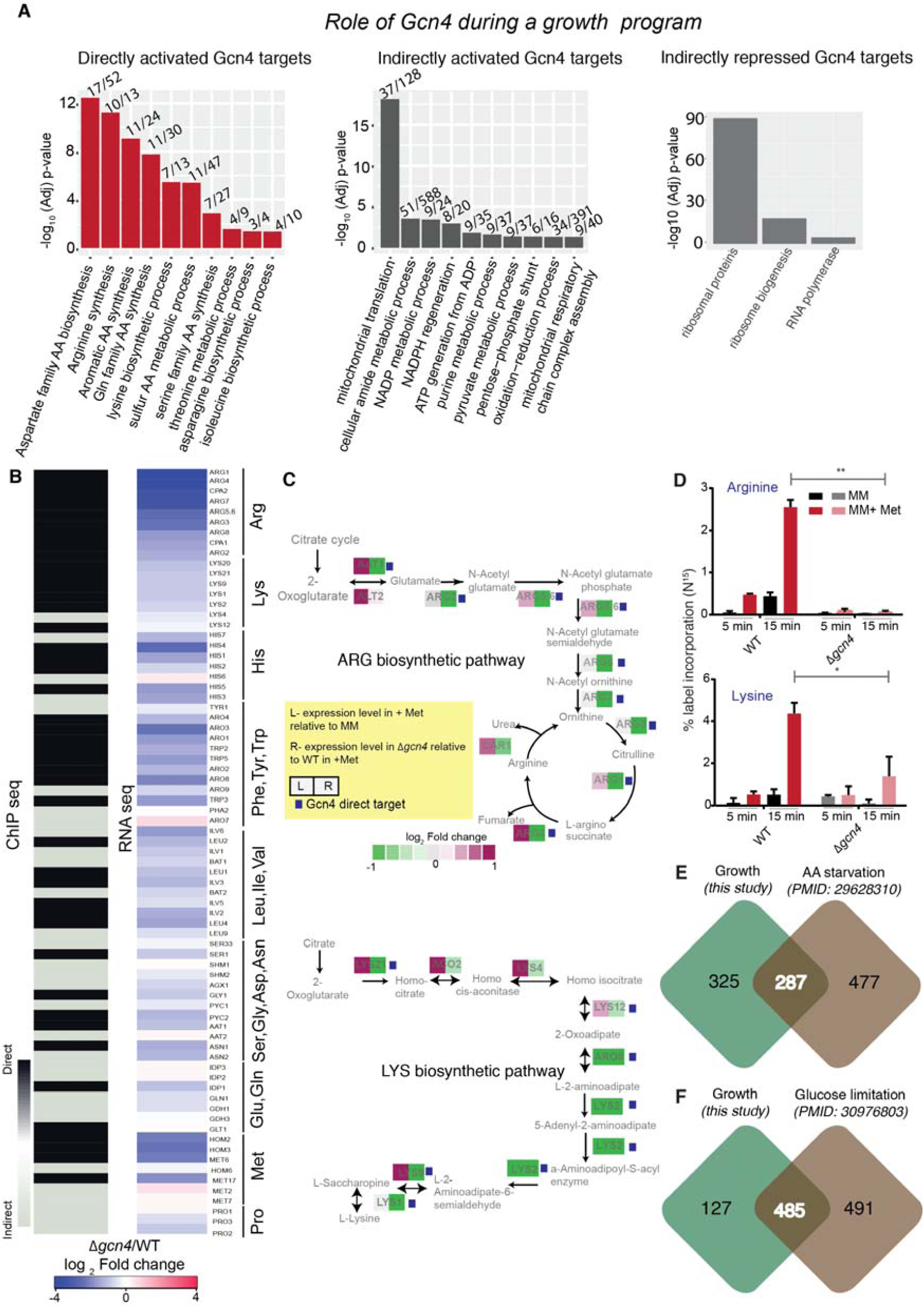
Direct and Indirect targets of Gcn4 during a growth program. A. Role of Gcn4 during a growth program (methionine addition): Bar plots shows enriched GO term and the corresponding -log10(p-value) for the genes which are directly or indirectly activated, or directly/indirectly repressed by Gcn4 when methionine is supplemented (growth program). Also see Supplementary Figure 7B. B. Comparing direct targets of Gcn4 regulon, and gene expression profiles of WT and *Δgcn4* cells (Gcn4 dependence) in a growth program. The heat map on the left shows whether the indicated gene (involved in amino acid biosynthesis) is directly or indirectly regulated by Gcn4 based on ChIP-Seq data from cells in MM+Met medium. The black color indicates a direct target of Gcn4 and grey indicates an indirect target (Gcn4 does not bind the promoter of this gene). The heatmap on the right shows the gene expression fold changes in *Δgcn4* relative to WT cells, grown in MM+Met medium. C. (Top panel) Representative pathway maps of the arginine biosynthetic pathway. This map shows the fold change in gene expression due to methionine (MM+Met compared to MM) in WT cells (left box), the change in gene expression due to loss of Gcn4 (WT compared to *Δgcn4*) in the presence of methionine (MM+Met) (right box). Genes that are direct targets of Gcn4 are also indicated with a small purple box next to the gene name. (Lower panel) A representative pathway map of the lysine biosynthetic pathway, represented similar to that of arginine biosynthesis. D. Increased arginine and lysine biosynthesis in a methionine dependent growth program depend entirely on Gcn4. Data from quantitative LC/MS/MS based metabolic flux analysis experiments, using N^15^ ammonium sulfate labeling to estimate new amino acid synthesis in a methionine and Gcn4 dependent manner, are shown. The comparisons are between WT and *Δgcn4* cells treated identically in MM+Met medium. The data are presented for arginine, lysine. Also see Figure Supplementary Figure 8. *p<0.05,**<0.01 (t-test). E. Overlap between potential Gcn4 binding targets identified by ChIP-seq in a growth program (this study), vs the targets identified under a severe amino acid starvation response [34]. F. Overlap between potential Gcn4 binding targets identified by ChIP-seq in a growth program (this study), vs the targets identified during moderate starvation induced by glucose limitation [52].

As discussed, in the presence of methionine Gcn4 directly upregulates the genes of various amino acid biosynthetic pathways (Figure 3A). In this context, subsets of amino acid biosynthetic genes are strikingly induced. Notably, every single gene of arginine biosynthetic pathway, and nearly every gene of lysine, histidine and branched chain amino acid biosynthetic pathways are directly activated by Gcn4 (Figure 3B and 3C, Supplementary Figure 7E). This suggests that Gcn4 might be critical for the supply of particularly arginine and lysine, during the methionine mediated anabolic program.

We estimated the functional contribution of methionine-induced Gcn4 towards individual amino acid biosynthesis, particularly that of arginine and lysine, using a targeted LC/MS/MS based approach [56] to measure amino acid synthesis flux, based on stable-isotope incorporation. Consistent with the transcriptome data, we found a strikingly increased amino acid biosynthesis in MM+Met, compared to MM, and expectedly; the loss of *GCN4* severely decreased the flux towards amino acid biosynthesis, including a near-complete loss of arginine and lysine biosynthesis (Figure 3D, Supplementary Figure 8). This reiterates that Gcn4 has a vital role in increasing the amino acid pools required during a methionine induced growth program, particularly regulating the synthesis of arginine and lysine.

How does the role of Gcn4 during this growth program compare to its role during extreme amino acid starvation? To understand this, we analyzed a publicly available ChIP seq data of Gcn4, where Gcn4 was induced during severe amino acid starvation (due to SM treatment) (34). Firstly, we compared potential Gcn4 targets, which are present 750bp around the Gcn4 peaks, identified in both the growth and the starvation condition. We found a 47% overlap between the Gcn4 targets during the methionine induced growth program, and under amino acid starvation [34] (Figure 3E, and Supplementary WS6) (Fisher’s Exact test P < 10^−10^). We also compared the targets identified in our study with a distinct, simpler starvation regime, where cells were only limited for glucose [52]. About 80% of the targets identified in this study overlap with that of the Gcn4 targets identified in the glucose limitation study [52] (Figure 3F, and Supplementary WS6) (Fisher’s Exact test P < 10^−10^). This indicates that the Gcn4 targets, particularly the activation of amino acid biosynthetic genes, are conserved irrespective of the growth status of the cell..

Finally, during starvation programs, Gcn4 negatively regulates (represses) ribosomal and translation related genes [31,34,35,44]. In agreement with these ChIP-seq studies in starvation conditions [34,35], we also find that Gcn4 indirectly represses translation related genes, except for the following-RPL14B, RPS9A, RPL36B and RRP5, these are directly repressed by Gcn4 under this condition (Supplementary Figure 7A and 7B). The distinction though is that when methionine is present, ribosomal genes are induced, but Gcn4 appears to temper the extent of this induction (as the loss of Gcn4 in this condition further increases ribosomal genes). Therefore, through this repressive activity, Gcn4 likely enables cells to manage the extent of ribosomal gene induction due to methionine.

To summarize, the role of Gcn4 in a methionine-dependent growth program can be broken into two parts. First, Gcn4 directly induces amino acid biosynthesis genes, as well as transamination reactions. As part of a feed-forward program, the nucleotide biosynthesis genes and the PPP (which complete the methionine-mediated anabolic program [20] are indirectly induced. Further, the ribosomal/translation related genes that are induced by methionine in wild-type cells are further induced upon the loss of Gcn4 in this condition, suggesting that Gcn4 manages the extent of ribosomal gene induction due to methionine. Notably, the core function of Gcn4, which is to increase amino acid (and nucleotide) synthesis, remains unchanged when cells are in a growth state or dealing with starvation. Importantly, Gcn4 is critical for the high rates of synthesis of arginine, lysine and histidine. However, the cellular outcome is different, because of this coincident activation of Gcn4 in conditions where ribosomal biogenesis and translation are high.

### Gcn4 globally represses arginine/lysine enriched genes, including the translational machinery

From our data thus far, it is clear that Gcn4 helps supply cells with several metabolites, particularly the amino acids arginine and lysine, when methionine triggers a growth program. Given this critical function of Gcn4 in arginine and lysine biosynthesis and supply, we wondered if there were correlations of lysine and arginine utilization in genes, and global gene expression programs controlled by Gcn4. Although the amino acid compositions of proteins are evolutionarily optimized, our understanding of amino acid supply vs demand remains woefully inadequate [57,58]. As amino acids are the building blocks of proteins, translation naturally depends on available amino acid pools in the cell. We therefore asked if there were categories of proteins that were particularly enriched for arginine and lysine, within the genome, and if this had any correlation with Gcn4 function. For this, we divided the total number of proteins in the *S. cerevisiae* genome into three bins based on the percentage of arginine and lysine content of the protein (%R+K). The bin1 comprises of 1491 proteins with the lowest percentage of R+K (bin1; < 10% R+K), bin2 has 3033 proteins with moderate %R+K content (bin2; 10-13% R+K), and bin3 comprises of the 1501 proteins, with very high %R+K (bin3; >13% R+K) (Figure 4A). We next asked if these bins were enriched for any groups of functional pathways (based on Gene Ontology). Bin1 and bin2 have very large, disparate groups of GO terms, with no unique enrichment. However, bin3 was significantly enriched for ribosomal and translation related genes (Figure 4B). This arginine and lysine distribution in translation related genes are significantly higher compared to genome wide distributions (Wilcox test, p-value < 10^−10^) (Figure 4C). Thus, translation related proteins are highly enriched for arginine and lysine amino acids.

**Figure 4:**
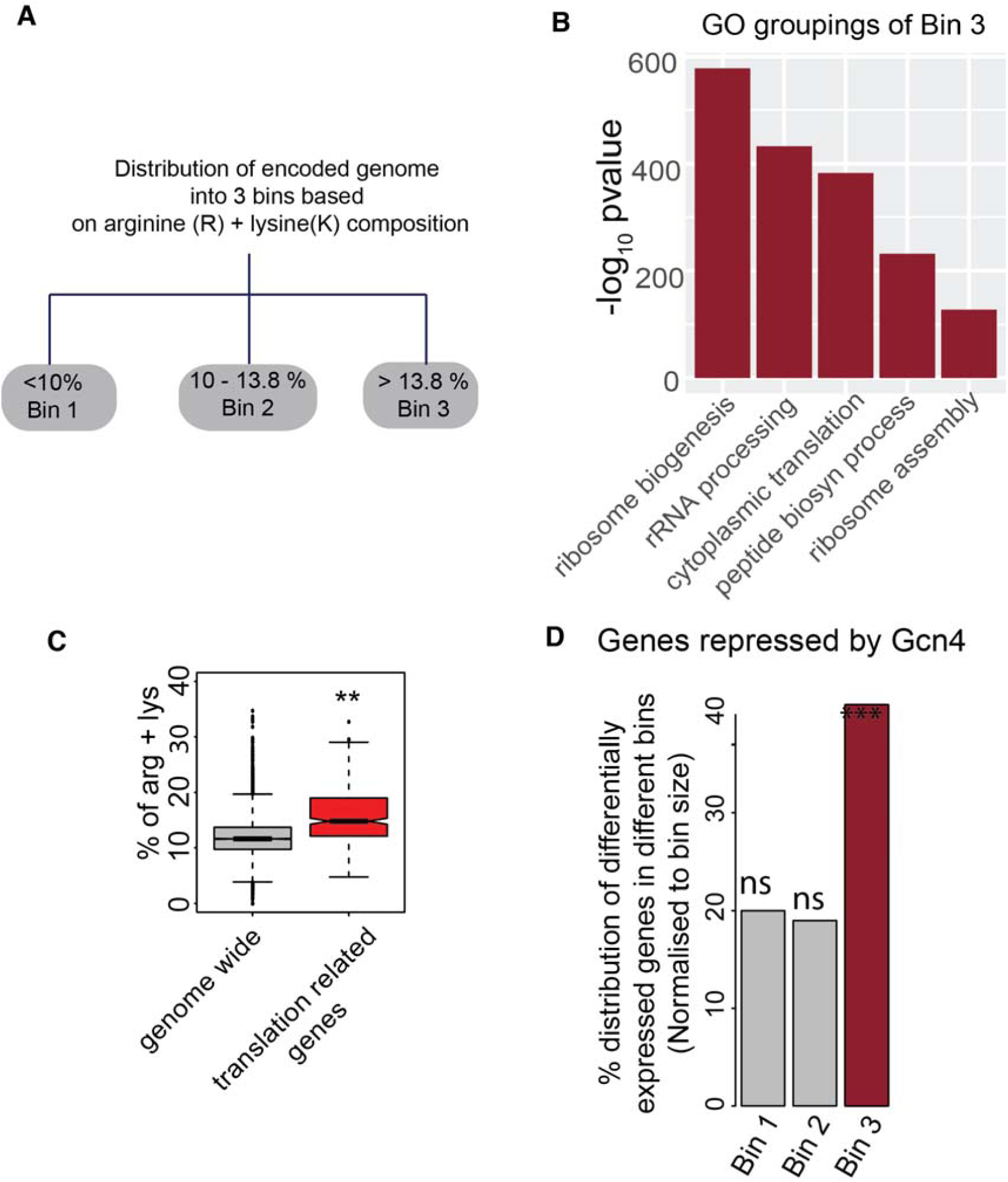
Gcn4 globally represses arginine/lysine enriched genes, including the translational machinery. A. “Binning” of the yeast proteome into three equal parts, based on the percentage of arginine and lysine in these proteins. The percentages of arginine and lysine (together) in these bins are indicated. B. GO based analysis reveals that bin3, which has the high percentage of arginine and lysine, is significantly enriched for ribosomal and translation related genes. The graph plots the most enriched GO term against -log10(P value). C. Boxplot, comparing the arginine and lysine composition of the entire proteome (excluding translation related genes), and the translation related genes. The translation related genes have a significantly higher than genome wide composition of arginine and lysine. D. Barplots, indicating in which bin (as shown in Figure 4A) the genes repressed by Gcn4 (i.e. induced in *Δgcn4*) fall under. A significant majority of the genes repressed by Gcn4 are enriched for arginine and lysine rich (Fisher exact test, p<10e-10).

Next, we asked if there is any correlation between the genes regulated by Gcn4 (in MM+Met), and the percentage of R+K encoded within these encoded proteins. Strikingly, we noticed that a very significant proportion of the genes that are repressed by Gcn4 fall in bin3 (∼40%, Fisher’s exact test P-value < 10^−10^) (Figure 4D & Supplementary WS5). Therefore, a significant proportion of the genes induced by methionine, and further induced in Δ*gcn4* are arginine and lysine rich. This suggests the possibility of a deeper management of overall, methionine-induced anabolism by Gcn4, where the translation of arginine and lysine enriched proteins will be required for high translation, and this requires Gcn4-dependent precursors.

### Gcn4 dependent outputs can sustain high translation capacity during growth

Given this striking observation, we asked if, in a growth program, cells could still sustain the synthesis of arginine and lysine rich genes if Gcn4 is absent. To evaluate this unambiguously, we designed inducible, luciferase-based reporters to estimate the translation of a several of R+K enriched genes, which are induced in cells by methionine (and further increased upon the loss of Gcn4) (Supplementary WS1). We designed a plasmid, in which the gene of interest (GOI; amplified from the genomic DNA of *S. cerevisiae*) was cloned in frame with the luciferase coding sequence, in such a way that the entire fragment (GOI+Luciferase) will be under the control of an inducible promoter (Supplementary Figure 9). Using this system, measuring luciferase activity after induction will estimate the specific translation of the specific arginine/lysine enriched gene, in any condition. This accounts for only newly synthesized protein, and therefore avoids mis-interpretations coming from already existing protein in the cells before methionine addition. We made reporters for 4 such candidate genes (RPL32, STM1, NHP2, RPS20) (Supplementary Figure 9).

First, we contextualized the expression of these lysine and arginine enriched genes (based on reporter activity) in wild-type cells, under either a growth or a starvation regime where Gcn4 expression is high. The conditions we compared were MM (low Gcn4 expression), addition of methionine (growth program, strong Gcn4 induction), and the addition of 3-AT (amino acid starvation condition, high Gcn4). In these conditions, we induced the reporters for Nhp2, Rpl32 and Rpl20 for 30 min, and compared luciferase activity. Here, the luciferase activity of all three reporters significantly increase in methionine supplemented conditions, and are decreased in the 3-AT condition (Figure 5A). This reiterates that the translational outcomes are entirely distinct in a growth or starvation program, despite high Gcn4 activity in both conditions.

**Figure 5:**
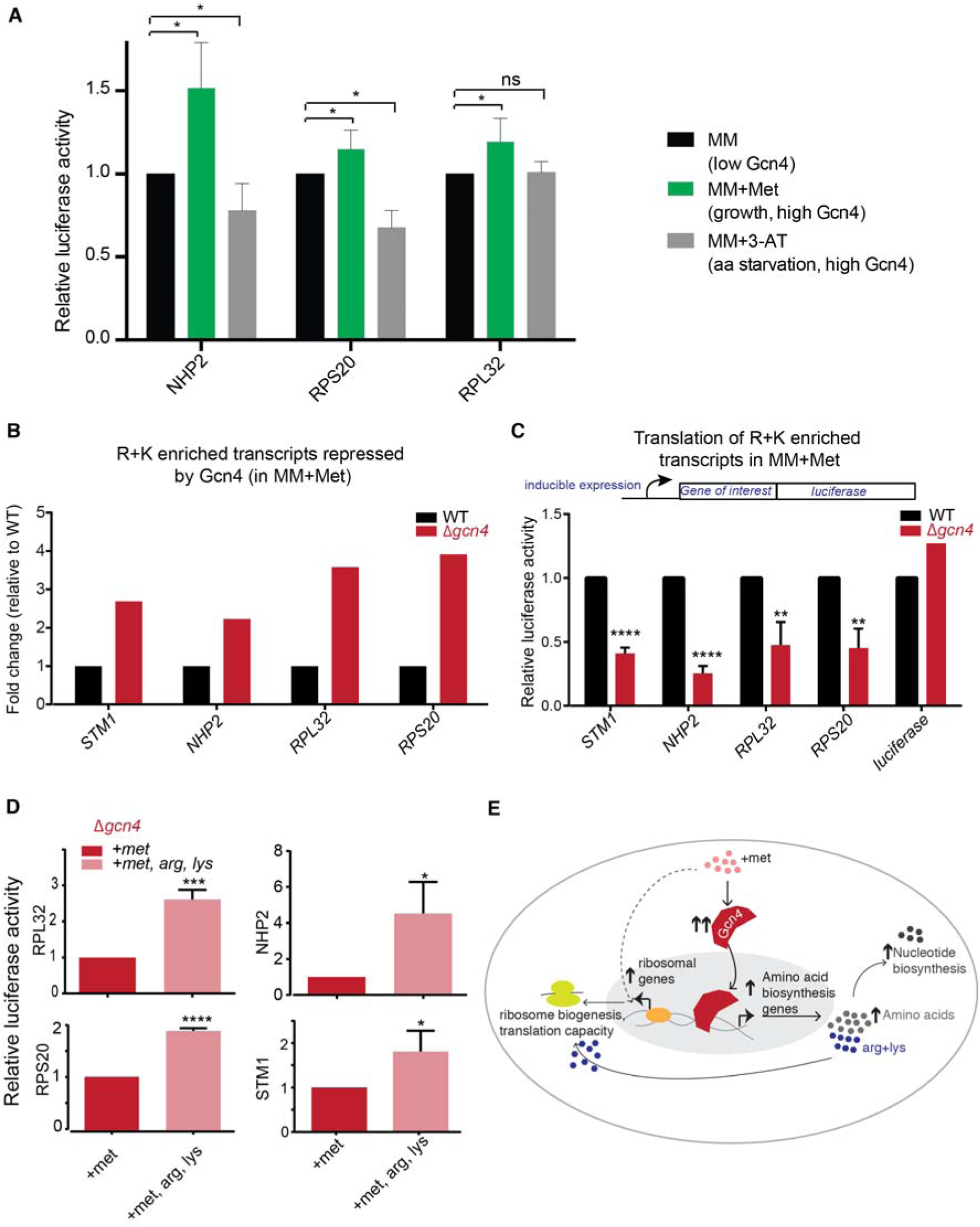
Gcn4 dependent outputs can sustain high translation capacity during growth. A. Translation of arg/lys genes increase during a growth program (methionine addition), and decrease during amino acid starvation (3-AT addition), both of which are conditions where Gcn4 is induced. B. Arg/lys enriched genes are induced in *Δgcn4* cells in methionine supplemented medium. Barplots comparing relative transcript amounts for selected, highly induced, arginine and lysine enriched genes, between WT and *Δgcn4* cells. Data shown are taken from the RNA seq data. C. Arg/lys enriched gene transcripts cannot be translated in *Δgcn4* cells in methionine supplemented medium. Barplots, comparing the relative amount of proteins translated (in a methionine-dependent manner), for the genes shown in Figure 5B. These selected genes were cloned in frame with luciferase in an inducible system, to create a translation-reporter, and translation of these were induced in WT or *Δgcn4* cells in methionine supplemented medium (MM+Met). Data shown are mean+/- SD from ≥ 3 biological replicates. *p<0.05,**p<0.01,****<0.0001 (t-test). D. Supplementing arginine and lysine restores translational capacity in *Δgcn4* cells. Barplots, comparing the relative amount of proteins translated (in a methionine-dependent manner), for the genes shown in Figure 5C, in methionine-supplemented medium (MM+Met) or in MM+Met+Arg+lys. Data shown are mean+/- SD from ≥ 3 biological replicates. *p<0.05,***p<0.001,****<0.0001 (t-test). E. A mechanistic model illustrating how high Gcn4 enables a methionine dependent anabolic response, by supplying amino acids, and maintaining translation capacity.

We now could specifically determine the importance of Gcn4 activity during a growth program (addition of methionine). First, we compared the extent of transcript expression for these lysine/arginine enriched transcripts, Nhp2, Rpl32, Rpl20 and Stm1, in wild-type and Δ*gcn4* cells in the presence of methionine (Figure 5B). The loss of Gcn4 in these conditions further increased expression of these transcripts, reiterating the role of Gcn4 as a (indirect) repressor of these genes. We next directly measured the translation of these genes, using the luciferase-based reporters of these genes. For this, using a similar experimental setup as earlier, we measured luciferase activity in wild-type and Δ*gcn4* cells after 30 minutes of induction with β-estradiol. Strikingly, all the candidate reporter genes showed a 3-5 fold reduction in translation in *GCN4* deficient cells, compared to WT cells (Figure 5C). These data reveal that Gcn4 is critically required to maintain the translation capacity of the cell, during this growth program. Finally, to determine if this reduced translation capacity in Δ*gcn4* is due to the reduced supply of arginine and lysine in these conditions, we carried out a rescue experiment with the addition of only these two amino acids. We supplied both amino acids (2mM each) to wild-type and Δ*gcn4* cells growing in the presence of methionine, and measured luciferase activity after induction. Notably, the supply of arginine and lysine substantially rescued the expression of these reporter proteins, by increasing their translation ∼2-3 fold (Figure 5D).

Collectively, we find that Gcn4 activity is central to sustain a growth program triggered by methionine. Gcn4 enables sufficient supply of amino acids, particularly arginine and lysine, for the translation of ribosomal proteins, while also tuning the amount of expression for these transcripts. This in turn maintains sufficient translational capacity needed by the cell, to sustain the anabolic program and drive growth. This is in contrast to a conventional starvation program due to amino acid limitation, where Gcn4 is also high. In such starvation contexts, the amounts of arginine and lysine enriched transcripts (including translation related transcripts) are low (and repressed by Gcn4), and the role of Gcn4 is in restoring amino acid levels.

## Discussion

A central theme emphasized in this study is mechanisms by which cells manage resource allocation, supply and demand during cell growth. Recent studies in model organisms like yeast and *E. coli* focus on protein translation, and the need to ‘buffer’ translation capacity during cell growth [8,59]. These studies alter our perception of how translation is regulated during high cell growth. However, the process of cell growth requires not just translation reserves (in the form of ribosomes), but also a constant supply of biosynthetic precursors to meet high demand. This includes: amino acids to sustain translation as well as drive metabolic functions, nucleotide synthesis (for DNA replication, transcription and ribosome biogenesis), and sufficient reductive capacity (for reductive biosynthesis). Even the production of ribosomes is an extremely resource-intensive process [60]. While our understanding of translation-regulation in these contexts is constantly improving, how metabolic and biosynthetic components are managed, and couple with translation, remain poorly understood.

Here, using yeast as a model, we obtain striking, mechanistic insight into how Gcn4 enables cells to sustain the supply of biosynthetic precursors, during a growth program (induced by methionine). In this growth program, methionine induces genes involved in ribosomal biogenesis and translation [20], which is the hallmark of a growth signature [27,60–63]. In addition, cells increase metabolic processes that sustain anabolism; primarily the pentose phosphate pathway, trans-amination reactions, an induction in amino acid biosynthesis, and nucleotide synthesis [20]. In particular, through this study, we now show how the Gcn4 transcription factor functions to critically support this growth program, by enabling high amino acid supply to maintain a sufficient translation capacity, as illustrated in a schematic model (Figure 5E).

Notably, we can now define the roles of Gcn4 during either a growth program, or a more conventionally studied starvation program. In a growth program (the methionine-induced context), genes required for ribosome biogenesis and translation that are induced have a nuanced regulation by Gcn4. Gcn4 represses ribosomal genes (consistent with earlier reports), and in this context thereby appears to balance or moderate the overall induction of ribosomal genes by methionine. The methionine-dependent induction of ribosomal genes is likely controlled by directly activating the TOR pathway [18,64–66], which is a regulator of ribosomal biogenesis. However, despite the high expression of ribosomal gene mRNAs in *Δgcn4* cells (in this growth program), and the indication of an apparent ‘growth signature’ transcriptional profile with high ribosomal transcripts, cells cannot sustain the required rates of protein synthesis, or maintain the high translation capacity required for growth. This is because the translation machinery itself is highly enriched for arginine and lysine amino acids, and so cannot be maintained at sufficient levels without a constant supply of lysine and arginine. In the presence of methionine, the increased synthesis (and therefore supply) of these two amino acids depends almost entirely on induced Gcn4. After all, to sustain high growth, and anabolic programs, cells need to maintain the required high rates of translation, and ribosomal capacity. Thus, through Gcn4, cells can deeply couple translation with metabolism, and manage sufficient resource allocations to sustain increased anabolism.

Notably, the specific transcriptional role of Gcn4 in growth or starvation programs remains conserved. Regardless of context, Gcn4 is required for amino acid biosynthesis (particularly lysine and arginine biosynthesis), and represses ribosomal genes. However, the different contexts completely alter the cellular outcomes, since in growth programs ribosomal genes are already high (and Gcn4 only tempers their expression), while in starvation programs ribosomal genes are low. The roles of Gcn4 have traditionally only been systematically examined during amino acid starvation or an ‘integrated stress response’ [29,67]. However, multiple studies now support a role for Gcn4 during contexts of high growth, including recent studies of the mammalian ortholog of Gcn4 (ATF4) which report high ATF4 activity in several cancers [36,37,68,69]. These studies suggest that ATF4 induction is critical for tumor progression during nutrient limitation, possibly by providing otherwise limiting metabolites [36,37], without clarity on the specific functions of ATF4 in these conditions. Separately, observations over decades note that many rapidly proliferating tumors depend on methionine [70,71], and methionine restriction critically determines tumor progression [23,24]. Here we directly demonstrate how Gcn4 provides biosynthetic precursor supply to sustain anabolism, in an otherwise limiting environment. Speculatively, could the ability of methionine to induce proliferation in cancers rest upon the induction of ATF4, which controls the supply of amino acids and other biosynthetic precursors?

Summarizing, here we address Gcn4 function during a growth program triggered by methionine. This expands the roles of a ‘starvation’ factor, during a contrasting, high anabolism state, showing how despite conserved function in both contexts, Gcn4 activity can lead to very different outcomes. Our study provides an illustrative perspective of how cells can manage the supply of important biosynthetic precursors with overall translation outputs, when a specific growth cues induce high biosynthetic demands that need to be coordinately sustained, in order to maintain anabolism and cell growth.

## Materials and Methods

### Strains and growth conditions

A fully prototropic yeast strain *Saccharomyces cerevisiae* strain from a CEN.PK background [72] was used in all experiments. For all the medium-shift experiments, overnight grown cultures were sub-cultured in fresh YPD (1% Yeast extract and 2% Peptone, 2% Glucose) medium with an initial OD_600_ of ∼0.2. Once the OD_600_ of the secondary culture reached 0.8 – 0.9, cells were pelleted down and washed and shifted to minimal media MM (MM-Yeast Nitrogen Base with glucose as a carbon source) and MM+Met (MM with 2mM methionine). For the luciferase assay (described later), overnight grown cultures were prepared by growing the cells in YPD with the antibiotic 1mM Nourseothricin (NAT). The secondary culture was started with an initial OD_600_ of ∼0.5 in YPD + NAT and incubated at 30°C and 250 rpm for 4 hours. After 4 hours of incubation, cells were washed once in MM, and shifted to MM +Met or MM+Met+arg+lys. 2mM concentration of each amino acid was used wherever required, unless mentioned otherwise. All the wash steps before shifting to minimal medium were done by centrifuging the cells at 3500 rpm for 90 seconds.

### Western blot analysis

Yeast cells with a 3x-FLAG epitope sequence chromosomally tagged at the carboxy-terminus of Gcn4 (endogenous locus) were used to quantify Gcn4 protein levels using western blotting (Supplementary table 1). Overnight grown cells were sub-cultured into fresh YPD medium, with an initial OD of 0.2 and grown to an OD_600_ of 0.8. Cells were pelleted down at 3500 rpm for 1.5 minutes, cell pellets were washed once in MM, re-harvested and shifted to MM and MM+ Met after 1 hour of the shift. ∼5 OD_600_ of cells were harvested by centrifugation, and proteins were precipitated in 400 µl of 10% trichloro acetic acid (TCA), and extracted by bead beating with glass beads. Lysates were centrifuged to precipitate all proteins, and total protein pellets were resuspended in 400 µl of SDS-Glycerol sample buffer. Protein concentrations were quantified using Bicinconinic assay kit (G-Biosciences, 786-570) and equal concentrations of proteins were loaded into the 4-12% Bis-tris polyacrylamide gel (Thermo Fisher,NP0322BOX) and resolved using electrophoresis. Resolved proteins were transferred to nitrocellulose membranes and detected by standard Western blotting using monoclonal anti-FLAG M2-mouse primary antibody (Sigma Aldrich, F1804) and HRP labelled anti-mouse IgG secondary antibody (Cell Signalling technology, 7076S). Blots were developed using enhanced chemiluminescence reagents (Advansta, K-12045) imaged using Image quant. A different part of each gel (cut out) was Coomassie stained in order to compare total protein loading amounts.

### mRNA sequencing and data analysis

Overnight grown cells of WT and *Δgcn4* strains were sub-cultured in YPD, with a starting OD_600_ of 0.2 and grown till they reached an OD_600_ of 0.8-0.9. YPD grown cells were pelleted down at 3500 rpm for 90 seconds and washed once with MM. Washed cells were shifted to MM and MM+Met, and cells remained in this fresh medium for ∼1 hr. The cells were collected an hour after this shift and RNA was isolated by a hot acid phenol method as described [73]. mRNA libraries were prepared using TruSeq RNA library preparation kit V2 (Illumina) and quality of the libraries were analyzed using bioanalyser (Agilent 2100) and libraries were sequenced for 51 cycles using Illumina HiSeq 2500 platform. For every experimental condition, data were obtained from two biological replicates. Normalized read counts between the biological replicates were well correlated (Figure S1). For each strain we obtained ∼30-35 million uniquely mapped reads. The raw data are available in NCBI-SRA under the accession PRJNA599001. The transcriptome data were aligned and mapped to the *S. cerevisiae S288C* genome downloaded from the saccharomyces genome database (SGD), using the Burrows-Wheeler Aligner [74] and the mapped reads with mapping quality of ≥ 20 were used for further analysis. Number of reads mapped to each gene was calculated and read count matrix was generated. The EdgeR package was used for normalization and differential gene expression analysis [75]. Differentially expressed genes with a fold change above 1.5 or 2 fold, with a stringent p-value cutoff of <= 0.0001 were considered for further analysis. Normalized read counts was calculated for every sample as described earlier [20]. Normalized read counts between the replicates are well correlated with the Pearson correlation coefficient (R) is more than 0.9 (Figure S1). GO analysis of the differentially expressed genes were carried out using g:Profiler [76].

### Chromatin Immunoprecipitation sequencing and data analysis

#### a. Cell growth conditions and sample collection

For ChIP sequencing, overnight grown cells were re-inoculated in fresh YPD medium (RM), with the initial OD_600_ of 0.2 and incubated at 30°C until the OD_600_ reached 0.8-0.9. Subsequently, 100 mL of culture was pelleted down, washed and shifted to MM and MM +Met. After 1 hour of the shift, cells were fixed using 1% formaldehyde, after which the fixing was quenched with 2.5M glycine.

#### b. Spheroplasting of fixed cells

Fixed cells were treated with 2-mercapto ethanol, and resuspended in 5 ml of spheroplasting buffer containing 1M sorbitol, 0.1M sodium citrate, 10mM EDTA, and distilled water, with 4mg/ml of lysing enzyme from *Trichoderma harzianum* (Sigma L1412-5G). This suspension was incubated at 37°C for 5 hours.

#### c. Lysis and ChIP

Spheroplasts were pelleted down at 1000 rpm, washed twice with Buffer 1 (0.25% Triton X100,10mM EDTA,0.5mM EGTA, 10mM sodium HEPES pH 6.5) and twice with Buffer 2 (200mM NaCl, 1mM EDTA, 0.5mM EGTA, 10mM Sodium HEPES pH 6.5), washed spheroplasts were resuspended in lysis buffer (50mM sodium HEPES pH 7.4, 1% Triton X, 140mM NaCl,0.1% Sodium deoxy cholate,10mM EDTA) and lysis and DNA fragmentation were carried out using a bioruptor (Diagenode, Nextgen) for 30 cycles (30 sec on and off cycles). Lysates were centrifuged to remove the debris and clear supernatant was used for chromatin immunoprecipitation (ChIP). Immunoprecipitation was carried out by incubating the lysate with the monoclonal anti-FLAG M2-mouse primary antibody (Sigma Aldrich, F1804) and protein G Dynabead (Invitrogen, 10004D). Beads were washed sequentially in low salt, high salt and LiCl buffers, TE buffer and protein-DNA complex were eluted using elution buffer as reported earlier [77]. Decrosslinking of the immuno-precipitated proteins were carried out by using a high concentration of NaCl and incubation at 65°C for 5 hours followed by proteinase-K treatment and DNA purification. Mock samples were also prepared in parallel, except the antibody treatment. Libraries were prepared for the purified IP DNA and mock samples (NEBNext Ultra II DNA library preparation kit, Catalog no-E7103L) and sequenced using Illumina platform HiSeq 2500. Two biological replicates were maintained for all the samples. The raw data are available in NCBI-SRA under the accession ID PRJNA599001.

ChIP sequencing reads were mapped to the *S. cerevisiae S288C* genome downloaded from SGD. The reads with mapping quality < 20 were discarded, and the remaining reads were used for further analysis. The number of reads mapped to every 100bp non-overlapping bins were calculated using ‘exomedepth’ function of R-package GenomicRanges [78]. Read counts were normalized by dividing the number of reads falling within each bin by the total number of reads fall within the range of µ±x, where, µ=mode of the distribution of read counts of each bin, x = median absolute deviation of all the bins that has a number of reads that are less than the mean of the distribution. Subsequently, the regions that have normalized read counts of above 2 were considered for further analysis. The binding regions which are separated by < 200bp were merged to give a single peak. The peaks which are conserved in both the replicates with the overlap of at least 50bp were considered as *bona fide* binding regions of Gcn4. Genes which are encoded around 750 bp on both sides of the peaks were listed in the Supplementary WS4.

### Peak feature annotation and motif analysis

Genomic features of the peaks were identified using the annotatePeak.pl function of the HOMER tool [48]. For motif analysis, nucleotide sequences corresponding to the peak intervals were extracted from the genome and motif identification was performed using ‘meme’ function of MEME-suite [49].

### Direct and Indirect target analysis

To annotate the genes corresponding to the peaks identified, the open reading frames that are encoded within 750 bp on both sides of the peak regions were considered as ‘possible Gcn4 binding targets’. Gene sets which are differentially expressed in *Δgcn4* relative to WT under MM+Met condition, with a fold change of > 2 (∼900 genes) were termed ‘Gcn4 regulatory targets’. While comparing these gene lists, the genes which intersect between these two gene sets are considered as ‘direct Gcn4 binding targets’ and the rest of the genes of ‘Gcn4 regulatory targets’ are ‘indirect targets of Gcn4’.

### Metabolic flux analysis using LC/MS/MS

To determine if the rates of biosynthesis of amino acids are altered in MM+Met and Gcn4 dependent manner, we measured ^15^N-label incorporation in amino acids. We used ^15^N-ammonium sulfate with all nitrogen atoms labelled. Cells grown in YPD were shifted to fresh minimal medium (with the appropriate carbon source as indicated), containing 0.5 X of unlabelled ammonium sulfate (0.25%) and MM+Met containing 0.5X of unlabelled ammonium sulfate (0.25%). After 1 hour of shift to minimal media, cells were pulsed with 0.25 % of ^15^N labelled ammonium sulfate and incubated for 5 minutes or 15 minutes as indicated. After the ^15^N pulse, metabolites were extracted and label incorporation into amino acids was analyzed using targeted LC/MS/MS protocols as described earlier [20,56]. Similarly, C^13^ labeled glucose was used to measure rate of biosynthesis of nucleotide.

### Luciferase based translation reporters for lysine and arginine enriched genes

To measure the translation of specific transcripts that encode arginine and lysine enriched proteins, the ORF of the following proteins, RPL32, NHP2, STM1, RPS20 were amplified from the genomic DNA isolated from WT CEN.PK strain of *S. cerevis*iae. The amplified ORFs (without the stop codon) were ligated to the luciferase cDNA amplified from pGL3 (Supplementary table 2). The resulting fragment with ‘*ORF*_*RK rich genes*_ *+ luciferase*’ were cloned in a centromeric (CEN.ARS) plasmid pSL207, a modified version of the plasmid used in the earlier study [79]. Luciferase expression in this construct is under the control of inducible promoter, which can be induced by ß-estradiol [79]. The resulting plasmids with the following genes RPL32, NHP2, STM1, RPS20 cloned in frame with luciferase and under the inducible GEV promoter were named pSL218, pSL221, pSL224, pSL234 respectively (Supplementary Figure 9 and Supplementary table 2). SL217 is a plasmid where only the luciferase cDNA amplified from pGL3 plasmid was cloned under the inducible promoter, serves as a control. All these plasmids generated have ampicillin selection (for amplification) and Nourseothricin resistant cassettes (NAT^r^) for selection in yeast. The generated plasmids were transformed to the WT and *Δgcn4* strains. To measure the translation of the genes cloned upstream of luciferase, these strains having plasmid were grown in YPD for overnight with an antibiotic NAT. Overnight grown cultures were shifted to fresh YPD+NAT with an initial OD_600_ of 0.4, and grown for 4 hours at 30°C. After 4 hrs of incubation in YPD, cells were washed and shifted to the MM and MM+Met. 75mM of 3-Amino triazole (3-AT) was used, wherever required. After 1 hour of the shift, cultures were split into two equal parts, one part of the culture was induced with 200nM ß-estradiol (Sigma Aldrich-E8875) and the other half was left as a mock-induced control. After 30 minutes of induction cells were harvested by centrifugation at 4°C at 3000 rpm and washed with lysis buffer (1X-PBS containing 1mM PMSF), after 3 washes, were resuspended in 200µl of lysis buffer. Lysed the cells by bead beating at 4° C (1min ON and 1 min OFF for 10min). After lysis, the protein concentrations were measured by BCA assay kit. Equal concentrations of protein were used for measuring the luciferase activity. Luciferase activity was measured using luciferase assay kit (Promega, E1500) and the activity was measured using a luminometer (Sirius, Titertek Berthold Detection systems). Luciferase activity (measured as Relative Light Units per Sec (RLU/sec)) were normalized with its respective uninduced control. Similar experiments were carried out under different media conditions supplemented with different amino acids, where the conditions are mentioned in the respective sections. The relative difference in luciferase activities between the strain types and media conditions were used to estimate changes in the active translation of these proteins.

### Statistical tests

R-Packages and Graph pad prism 7 were used for visualizing data and performing statistical tests. The respective statistical tests used, and the statistical significance was mentioned wherever required.

## Acknowledgements

We acknowledge extensive use of the NCBS/inStem/CCAMP next generation sequencing facilities, and the NCBS/inStem/CCAMP mass spectrometry facilities. RS acknowledges support from SERB National Postdoctoral Fellowship (PDF/2016/001877), DST, Govt. of India. SL acknowledges support from a DBT-Wellcome trust India Alliance Intermediate Fellowship (IA/I/14/2/501523), and intramural support for this study. AS acknowledges support from a DBT-Wellcome trust India Alliance Intermediate Fellowship (IA/I/16/2/502711).

## Author contributions

RS and SL conceived the study, RS, ASW, ASN and SL designed experiments, RS and AW performed experiments, RS, ASW, AS and SL analyzed data, RS, AS and SL wrote the manuscript.

## Supporting Information and Legends

The Supporting Information is provided as the following files:

1. Supplementary results, Supplementary Figures 1-9, and Supplementary Tables 1 and 2 (single pdf file)
2. Supplementary Worksheet 1 (.xlsx format), of differentially expressed genes in the indicated conditions.
3. Supplementary Worksheet 2 (.xlsx format), list of anabolic and translation related genes induced in the indicated conditions.
4. Supplementary Worksheet 3 (.xlsx format), of GO categories of genes up/down regulated in the indicated conditions.
5. Supplementary Worksheet 4 (.xlsx format), of Gcn4 targets based on ChIP-seq analysis.
6. Supplementary Worksheet 5 (.xlsx format), of genes repressed by Gcn4 that are in bin 3 (from Figure 4).
7. Supplementary Worksheet 6 (.xlsx format), of genes induced by Gcn4 and repressed by Gcn4, and overlap of Gcn4 targets from starvation studies.

## References

1. Gresham D, Boer VM, Caudy A, Ziv N, Brandt NJ, Storey JD, et al. System-level analysis of genes and functions affecting survival during nutrient starvation in Saccharomyces cerevisiae. Genetics. 2011; doi: 10.1534/genetics.110.120766

2. Boer VMVM, Crutchfield CACA, Bradley PHPH, Botstein D, Rabinowitz JDJD. Growth-limiting Intracellular Metabolites in Yeast Growing under Diverse Nutrient Limitations. Mol Biol Cell. 2010;21: 198–211. doi: 10.1091/mbc.E09-07-0597

3. Saldanha A, Brauer M, Botstein D. Nutritional Homeostasis in Batch and Steady-State Culture of Yeast. Mol Biol Cell. 2004;15: 4089–104. doi: 10.1091/mbc.E04-04-0306

4. Dikicioglu D, Karabekmez E, Rash B, Pir P, Kirdar B, Oliver SG. How yeast re-programmes its transcriptional profile in response to different nutrient impulses. BMC Syst Biol. BioMed Central; 2011;5: 148. doi: 10.1186/1752-0509-5-148

5. Gutteridge A, Pir P, Castrillo JI, Charles PD, Lilley KS, Oliver SG. Nutrient control of eukaryote cell growth: a systems biology study in yeast. BMC Biol. BioMed Central; 2010;8: 68. doi: 10.1186/1741-7007-8-68

6. Brauer MJ, Yuan J, Bennett BD, Lu W, Kimball E, Botstein D, et al. Conservation of the metabolomic response to starvation across two divergent microbes. Proc Natl Acad Sci. 2006;103: 19302–19307.

7. Gurvich Y, Leshkowitz D, Barkai N. Dual role of starvation signaling in promoting growth and recovery. PLoS Biol. 2017;15: 1–28. doi: 10.1371/journal.pbio.2002039

8. Metzl-Raz E, Kafri M, Yaakov G, Soifer I, Gurvich Y, Barkai N. Principles of cellular resource allocation revealed by condition-dependent proteome profiling. Elife. eLife Sciences Publications, Ltd; 2017;6: e28034. doi: 10.7554/eLife.28034

9. Zaman S, Lippman SI, Zhao X, Broach JR. How Saccharomyces Responds to Nutrients. Annu Rev Genet. 2008;42: 27–81. doi: 10.1146/annurev.genet.41.110306.130206

10. Cai L, Sutter BM, Li B, Tu BP. Acetyl-CoA induces cell growth and proliferation by promoting the acetylation of histones at growth genes. Mol Cell. Elsevier Inc.; 2011;42: 426–37. doi: 10.1016/j.molcel.2011.05.004

11. Cai L, Tu BP. On acetyl-CoA as a gauge of cellular metabolic state. Cold Spring Harb Perspect Biol. 2011;76: 195–202.

12. Kuang Z, Cai L, Zhang X, Ji H, Tu BP, Boeke JD. High-temporal-resolution view of transcription and chromatin states across distinct metabolic states in budding yeast. Nat Struct Mol Biol. 2014/08/31. 2014;21: 854–863. doi: 10.1038/nsmb.2881

13. Shi L, Tu BPBP. Acetyl-CoA induces transcription of the key G1 cyclin CLN3 to promote entry into the cell division cycle in Saccharomyces cerevisiae. Proc Natl Acad Sci. 2013;110: 7318–7323. doi: 10.1073/pnas.1302490110

14. Krishna S, Laxman S. A minimal “push – pull “bistability model explains oscillations between quiescent and proliferative cell states. Lew DJ, editor. Mol Biol Cell. 2018;29: 2243–2258. doi: 10.1091/mbc.E18-01-0017

15. Wellen KE, Thompson CB. Cellular metabolic stress: considering how cells respond to nutrient excess. Mol Cell. Elsevier Inc.; 2010;40: 323–32. doi: 10.1016/j.molcel.2010.10.004

16. Rowicka M, Kudlicki A, Tu BP, Otwinowski Z. High-resolution timing of cell cycle-regulated gene expression. Proc Natl Acad Sci U S A. 2007;104: 16892–7. doi: 10.1073/pnas.0706022104

17. Pedro MB, Madeo F, Pietrocola F, Galluzzi L, Bravo-San Pedro JM, Madeo F, et al. Review Acetyl Coenzyme AC: A Central Metabolite and Second Messenger. Cell Metab. United States; 2015;21: 805–821. doi: 10.1016/j.cmet.2015.05.014

18. Sutter BM, Wu X, Laxman S, Tu BP. Methionine inhibits autophagy and promotes growth by inducing the SAM-responsive methylation of PP2A. Cell. 2013;154: 403–15. doi: 10.1016/j.cell.2013.06.041

19. Lees EK, Banks R, Cook C, Hill S, Morrice N, Grant L, et al. Direct comparison of methionine restriction with leucine restriction on the metabolic health of C57BL/6J mice. Sci Rep. Nature Publishing Group UK; 2017;7: 9977. doi: 10.1038/s41598-017-10381-3

20. Walvekar AS, Srinivasan R, Gupta R, Laxman S. Methionine coordinates a hierarchically organized anabolic program enabling proliferation. Mol Biol Cell. American Society for Cell Biology (mboc); 2018;29: 3183–3200. doi: 10.1091/mbc.E18-08-0515

21. Halpern BC, Clark BR, Hardy DN, Halpern RM, Smith RA. The effect of replacement of methionine by homocystine on survival of malignant and normal adult mammalian cells in culture. Proc Natl Acad Sci U S A. 1974;71: 1133–6. doi: 10.1073/pnas.71.4.1133

22. Sugimura T, Birnbaum SM, Winitz M, Greenstein JP. Quantitative nutritional studies with water-soluble, chemically defined diets. VIII. The forced feeding of diets each lacking in one essential amino acid. Arch Biochem Biophys. 1959;81: 448–455. doi: 10.1016/0003-9861(59)90225-5

23. Gao X, Sanderson SM, Dai Z, Reid MA, Cooper DE, Lu M, et al. Dietary methionine restriction targets one carbon metabolism in humans and produces broad therapeutic responses in cancer. bioRxiv. 2019; 627364. doi: 10.1101/627364

24. Gao X, Sanderson SM, Dai Z, Reid MA, Cooper DE, Lu M, et al. Dietary methionine influences therapy in mouse cancer models and alters human metabolism. Nature. 2019/07/31. England; 2019;572: 397–401. doi: 10.1038/s41586-019-1437-3

25. Wu X, Tu BP. Selective regulation of autophagy by the Iml1-Npr2-Npr3 complex in the absence of nitrogen starvation. Mol Biol Cell. 2011;22: 4124–4133. doi: 10.1091/mbc.E11-06-0525

26. Laxman S, Sutter B, Tu BP. Methionine is a signal of amino acid sufficiency that inhibits autophagy through the methylation of PP2A. Autophagy. 2014;10: 386–387. doi: 10.4161/auto.27485

27. Brauer MJMJ, Huttenhower C, Airoldi EMEM, Rosenstein R, Matese JCJC, Gresham D, et al. Coordination of Growth Rate, Cell Cycle, Stress Response, and Metabolic Activity in Yeast. Mol Biol Cell. 2008;19: 352–267. doi: 10.1091/mbc.E07-08-0779

28. Hinnebusch AG. Translational Regulation Of GCN4 and the General Amino Acid Control of yeast. Annu Rev Microbiol. 2005;59: 407–50. doi: 10.1146/annurev.micro.59.031805.133833

29. Mascarenhas C, Edwards-Ingram LC, Zeef L, Shenton D, Ashe MP, Grant CM. Gcn4 is required for the response to peroxide stress in the yeast Saccharomyces cerevisiae. Mol Biol Cell. 2008/04/16. The American Society for Cell Biology; 2008;19: 2995–3007. doi: 10.1091/mbc.e07-11-1173

30. Yang R, Wek SA, Wek RC. Glucose limitation induces GCN4 translation by activation of Gcn2 protein kinase. Mol Cell Biol. American Society for Microbiology; 2000;20: 2706–2717. doi: 10.1128/mcb.20.8.2706-2717.2000

31. Natarajan K, Meyer MR, Jackson BM, Slade D, Roberts C, Hinnebusch AG, et al. Transcriptional profiling shows that Gcn4p is a master regulator of gene expression during amino acid starvation in yeast. Mol Cell Biol. 2001;21: 4347–68. doi: 10.1128/MCB.21.13.4347-4368.2001

32. Hinnebusch AG, Asano K, Olsen DS, Phan L, Nielsen KH, Valásek L. Study of translational control of eukaryotic gene expression using yeast. Ann N Y Acad Sci. United States; 2004;1038: 60–74. doi: 10.1196/annals.1315.012

33. Hinnebusch AG. Translational control of GCN4: an in vivo barometer of initiation-factor activity. Trends Biochem Sci. England; 1994;19: 409–414. doi: 10.1016/0968-0004(94)90089-2

34. Rawal Y, Chereji R V, Valabhoju V, Qiu H, Ocampo J, Clark DJ, et al. Gcn4 Binding in Coding Regions Can Activate Internal and Canonical 5’ Promoters in Yeast. Mol Cell. 2018/04/05. 2018;70: 297-311.e4. doi: 10.1016/j.molcel.2018.03.007

35. Mittal N, Guimaraes JC, Gross T, Schmidt A, Vina-Vilaseca A, Nedialkova DD, et al. The Gcn4 transcription factor reduces protein synthesis capacity and extends yeast lifespan. Nat Commun. Nature Publishing Group UK; 2017;8: 457. doi: 10.1038/s41467-017-00539-y

36. Ye J, Kumanova M, Hart LS, Sloane K, Zhang H, De Panis DN, et al. The GCN2-ATF4 pathway is critical for tumour cell survival and proliferation in response to nutrient deprivation. EMBO J. 2010;29: 2082–96. doi: 10.1038/emboj.2010.81

37. Tameire F, Verginadis II, Leli NM, Polte C, Conn CS, Ojha R, et al. ATF4 couples MYC-dependent translational activity to bioenergetic demands during tumour progression. Nat Cell Biol. 2019/07/01. Springer US; 2019;21: 889–899. doi: 10.1038/s41556-019-0347-9

38. Xiao L, Grove A. Coordination of Ribosomal Protein and Ribosomal RNA Gene Expression in Response to TOR Signaling. Curr Genomics. Bentham Science Publishers Ltd; 2009;10: 198–205. doi: 10.2174/138920209788185261

39. Airoldi EM, Huttenhower C, Gresham D, Lu C, Caudy AA, Dunham MJ, et al. Predicting cellular growth from gene expression signatures. PLoS Comput Biol. 2009/01/02. Public Library of Science; 2009;5: e1000257–e1000257. doi: 10.1371/journal.pcbi.1000257

40. Hinnebusch AG, Natarajan K. Gcn4p, a master regulator of gene expression, is controlled at multiple levels by diverse signals of starvation and stress. Eukaryot Cell. American Society for Microbiology; 2002;1: 22–32. doi: 10.1128/ec.01.1.22-32.2002

41. Pakos-Zebrucka K, Koryga I, Mnich K, Ljujic M, Samali A, Gorman AM. The integrated stress response. EMBO Rep. 2016/09/14. John Wiley and Sons Inc.; 2016;17: 1374–1395. doi: 10.15252/embr.201642195

42. Akhter A, Rosonina E. Chromatin Association of Gcn4 Is Limited by Post-translational Modifications Triggered by its DNA-Binding in Saccharomyces cerevisiae. Genetics. 2016/10/21. Genetics Society of America; 2016;204: 1433–1445. doi: 10.1534/genetics.116.194134

43. Albrecht G, Mösch HU, Hoffmann B, Reusser U, Braus GH. Monitoring the Gcn4 protein-mediated response in the yeast Saccharomyces cerevisiae. J Biol Chem. United States; 1998;273: 12696–12702. doi: 10.1074/jbc.273.21.12696

44. Joo YJ, Kim J, Kang U, Yu M, Kim J. Gcn4p-mediated transcriptional repression of ribosomal protein genes under amino-acid starvation. EMBO J. Nature Publishing Group; 2010;30: 859–872. doi: 10.1038/emboj.2010.332

45. Bose T, Lee KK, Lu S, Xu B, Harris B, Slaughter B, et al. Cohesin proteins promote ribosomal RNA production and protein translation in yeast and human cells. PLoS Genet. 2012/06/14. Public Library of Science; 2012;8: e1002749–e1002749. doi: 10.1371/journal.pgen.1002749

46. Saint M, Sawhney S, Sinha I, Singh RP, Dahiya R, Thakur A, et al. The TAF9 C-terminal conserved region domain is required for SAGA and TFIID promoter occupancy to promote transcriptional activation. Mol Cell Biol. 2014/02/18. American Society for Microbiology; 2014;34: 1547–1563. doi: 10.1128/MCB.01060-13

47. McMillan J, Lu Z, Rodriguez JS, Ahn T-H, Lin Z. YeasTSS: an integrative web database of yeast transcription start sites. Database (Oxford). Oxford University Press; 2019;2019: baz048. doi: 10.1093/database/baz048

48. Heinz S, Benner C, Spann N, Bertolino E, Lin YC, Laslo P, et al. Simple combinations of lineage-determining transcription factors prime cis-regulatory elements required for macrophage and B cell identities. Mol Cell. 2010;38: 576–589. doi: 10.1016/j.molcel.2010.05.004

49. Bailey TL, Boden M, Buske FA, Frith M, Grant CE, Clementi L, et al. MEME SUITE: tools for motif discovery and searching. Nucleic Acids Res. 2009/05/20. Oxford University Press; 2009;37: W202–W208. doi: 10.1093/nar/gkp335

50. Arndt K, Fink GR. GCN4 protein, a positive transcription factor in yeast, binds general control promoters at all 5’ TGACTC 3’ sequences. Proc Natl Acad Sci U S A. 1986;83: 8516–8520. doi: 10.1073/pnas.83.22.8516

51. Oakley MG, Dervan PB. Structural motif of the GCN4 DNA binding domain characterized by affinity cleaving. Science. United States; 1990;248: 847–850. doi: 10.1126/science.2111578

52. Holland P, Bergenholm D, Börlin CS, Liu G, Nielsen J. Predictive models of eukaryotic transcriptional regulation reveals changes in transcription factor roles and promoter usage between metabolic conditions. Nucleic Acids Res. Oxford University Press; 2019;47: 4986–5000. doi: 10.1093/nar/gkz253

53. Aow JSZ, Xue X, Run J-Q, Lim GFS, Goh WS, Clarke ND. Differential binding of the related transcription factors Pho4 and Cbf1 can tune the sensitivity of promoters to different levels of an induction signal. Nucleic Acids Res. 2013/04/04. Oxford University Press; 2013;41: 4877–4887. doi: 10.1093/nar/gkt210

54. Kribelbauer JF, Rastogi C, Bussemaker HJ, Mann RS. Low-Affinity Binding Sites and the Transcription Factor Specificity Paradox in Eukaryotes. Annu Rev Cell Dev Biol. 2019/07/05. 2019;35: 357–379. doi: 10.1146/annurev-cellbio-100617-062719

55. Nishizawa M, Komai T, Morohashi N, Shimizu M, Toh-e A. Transcriptional repression by the Pho4 transcription factor controls the timing of SNZ1 expression. Eukaryot Cell. 2008/04/11. American Society for Microbiology (ASM); 2008;7: 949–957. doi: 10.1128/EC.00366-07

56. Walvekar A, Rashida Z, Maddali H, Laxman S. A versatile LC-MS/MS approach for comprehensive, quantitative analysis of central metabolic pathways. Wellcome open Res. F1000 Research Limited; 2018;3: 122. doi: 10.12688/wellcomeopenres.14832.1

57. Hofmeyr JHS, Cornish-Bowden A. Regulating the cellular economy of supply and demand. FEBS Lett. 2000;476: 47–51. doi: 10.1016/S0014-5793(00)01668-9

58. Hofmeyr J-HSHS. The harmony of the cell: the regulatory design of cellular processes. Wolkenhauer O, Wellstead P, Cho K-H, editors. Essays Biochem. 2008;45: 57–66. doi: 10.1042/bse0450057

59. Hui S, Silverman JM, Chen SS, Erickson DW, Basan M, Wang J, et al. Quantitative proteomic analysis reveals a simple strategy of global resource allocation in bacteria. Mol Syst Biol. BlackWell Publishing Ltd; 2015;11: 784. doi: 10.15252/msb.20145697

60. Warner JR, Vilardell J, Sohn JH. Economics of ribosome biosynthesis. Cold Spring Harb Symp Quant Biol. 2001;66: 567–74. doi: 10.1101/sqb.2001.66.567

61. Bosdriesz E, Molenaar D, Teusink B, Bruggeman FJ. How fast-growing bacteria robustly tune their ribosome concentration to approximate growth-rate maximization. FEBS J. 2015/03/26. John Wiley & Sons, Ltd; 2015;282: 2029–2044. doi: 10.1111/febs.13258

62. Klumpp S, Scott M, Pedersen S, Hwa T. Molecular crowding limits translation and cell growth. Proc Natl Acad Sci U S A. 2013/09/30. National Academy of Sciences; 2013;110: 16754–16759. doi: 10.1073/pnas.1310377110

63. Tu BP, Tu BP, Kudlicki A, Rowicka M, Mcknight SL. Logic of the Yeast Metabolic CycleC: of Cellular Processes. Science. 2005; doi: 10.1126/science.1120499

64. Martin DE, Soulard A, Hall MN. TOR regulates ribosomal protein gene expression via PKA and the Forkhead transcription factor FHL1. Cell. 2004;119: 969–79. doi: 10.1016/j.cell.2004.11.047

65. Mayer C, Grummt I. MRibosome biogenesis and cell growth: mTOR coordinates transcription by all three classes of nuclear RNA polymerases. Oncogene. England; 2006;25: 6384–6391. doi: 10.1038/sj.onc.1209883

66. Gu X, Orozco JM, Saxton RA, Condon KJ, Liu GY, Krawczyk PA, et al. SAMTOR is an S - adenosylmethionine sensor for the mTORC1 pathway. Science (80-). 2017;818: 813–818.

67. Dey S, Baird TD, Zhou D, Palam LR, Spandau DF, Wek RC. Both transcriptional regulation and translational control of ATF4 are central to the integrated stress response. J Biol Chem. 2010/08/23. American Society for Biochemistry and Molecular Biology; 2010;285: 33165–33174. doi: 10.1074/jbc.M110.167213

68. Singleton DC, Harris AL. Targeting the ATF4 pathway in cancer therapy. Expert Opin Ther Targets. 2012/09/26. England; 2012;16: 1189–1202. doi: 10.1517/14728222.2012.728207

69. Pällmann N, Livgård M, Tesikova M, Zeynep Nenseth H, Akkus E, Sikkeland J, et al. Regulation of the unfolded protein response through ATF4 and FAM129A in prostate cancer. Oncogene. 2019/07/16. England; 2019;38: 6301–6318. doi: 10.1038/s41388-019-0879-2

70. Breillout F, Antoine E, Poupon MF. Methionine dependency of malignant tumors: a possible approach for therapy. J Natl Cancer Inst. 1990;82: 1628–32. doi: 10.1093/jnci/82.20.1628

71. Poirson-Bichat F, Gonçalves RA, Miccoli L, Dutrillaux B, Poupon MF. Methionine depletion enhances the antitumoral efficacy of cytotoxic agents in drug-resistant human tumor xenografts. Clin Cancer Res. 2000;6: 643–53.

72. van Dijken JP, Bauer, Brambilla, Duboc, Francois, Gancedo, et al. An interlaboratory comparison of physiological and genetic properties of four Saccharomyces cerevisiae strains. Enzyme Microb Technol. 2000;26: 706–714. doi: 10.1016/S0141-0229(00)00162-9

73. Collart MA, Oliviero S. Preparation of Yeast RNA. Current Protocols in Molecular Biology. Hoboken, NJ, USA: John Wiley & Sons, Inc.; 2001. doi: 10.1002/0471142727.mb1312s23

74. Li H, Durbin R. Fast and accurate short read alignment with Burrows-Wheeler transform. Bioinformatics. 2009;25: 1754–1760. doi: 10.1093/bioinformatics/btp324

75. Robinson MD, McCarthy DJ, Smyth GK. edgeR: a Bioconductor package for differential expression analysis of digital gene expression data. Bioinformatics. 2010;26: 139–40. doi: 10.1093/bioinformatics/btp616

76. Raudvere U, Kolberg L, Kuzmin I, Arak T, Adler P, Peterson H, et al. g:Profiler: a web server for functional enrichment analysis and conversions of gene lists (2019 update). Nucleic Acids Res. Oxford University Press; 2019;47: W191–W198. doi: 10.1093/nar/gkz369

77. Lelandais G, Blugeon C, Merhej J. ChIPseq in Yeast Species: From Chromatin Immunoprecipitation to High-Throughput Sequencing and Bioinformatics Data Analyses. Methods Mol Biol. United States; 2016;1361: 185–202. doi: 10.1007/978-1-4939-3079-1_11

78. Lawrence M, Huber W, Pagès H, Aboyoun P, Carlson M, Gentleman R, et al. Software for computing and annotating genomic ranges. PLoS Comput Biol. 2013/08/08. Public Library of Science; 2013;9: e1003118–e1003118. doi: 10.1371/journal.pcbi.1003118

79. McIsaac RS, Silverman SJ, McClean MN, Gibney PA, Macinskas J, Hickman MJ, et al. Fast-acting and nearly gratuitous induction of gene expression and protein depletion in Saccharomyces cerevisiae. Mol Biol Cell. 2011/09/30. The American Society for Cell Biology; 2011;22: 4447–4459. doi: 10.1091/mbc.E11-05-0466

