## Supplementary figures and tables for "Genome-scale reconstruction of Gcn4/ATF4 networks driving a growth program" (from another version of this preprint: version 1, 2020-01-29)

**Supplementary Figures and legends.**

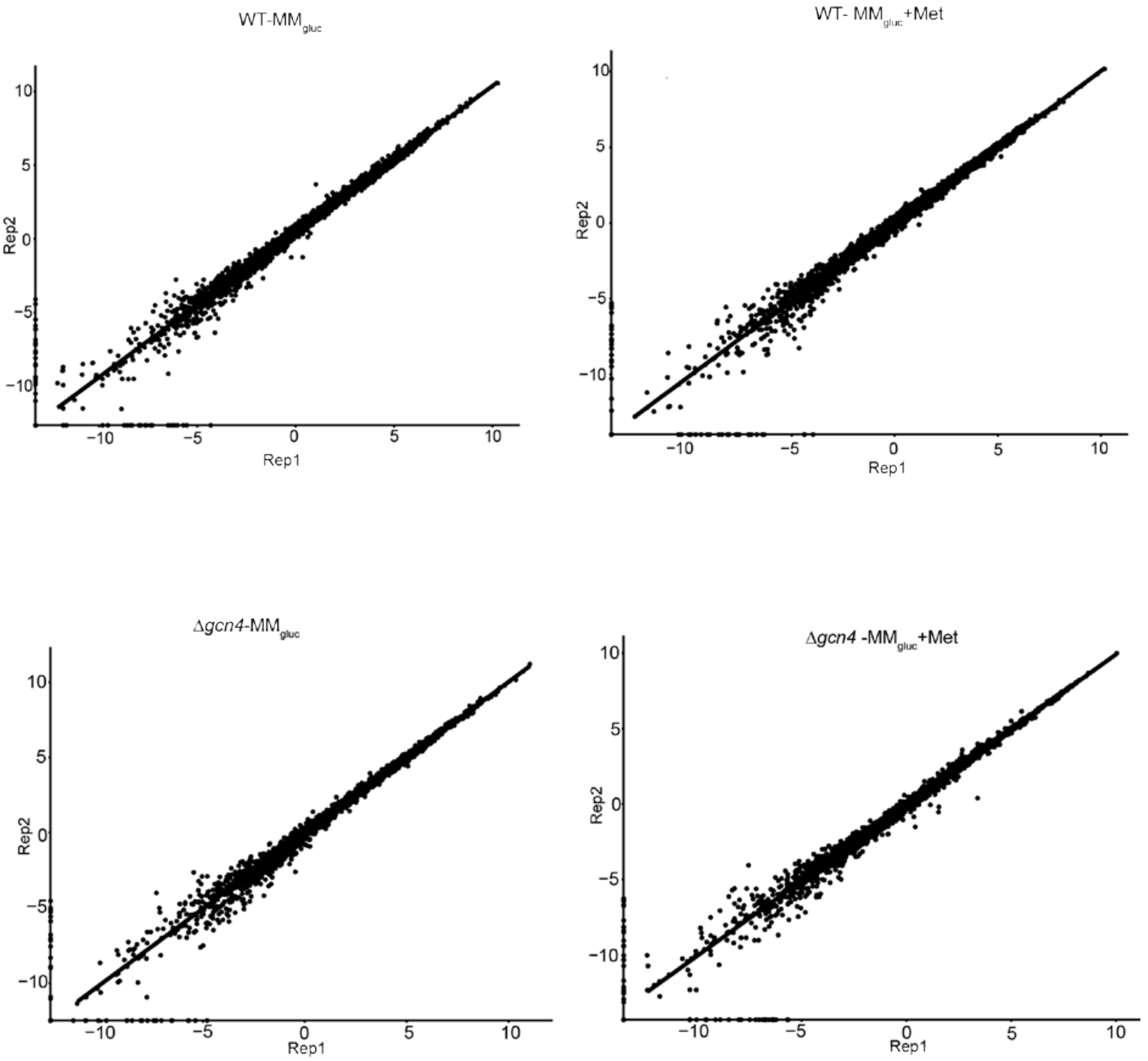


**Figure S1: RNA-seq correlation plots**
Correlation plots between replicates of the RNA sequencing data, from both WT and Δ*gcn4* cells, for the indicated media conditions. Normalised read counts of the RNA sequencing data are plotted. Replicates shows good correlation with a Pearson correlation coefficient, R = 0.9.

**A**

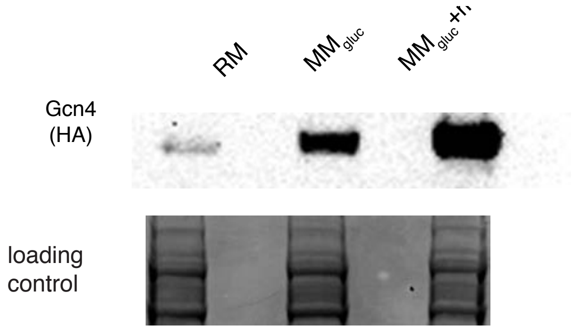


B

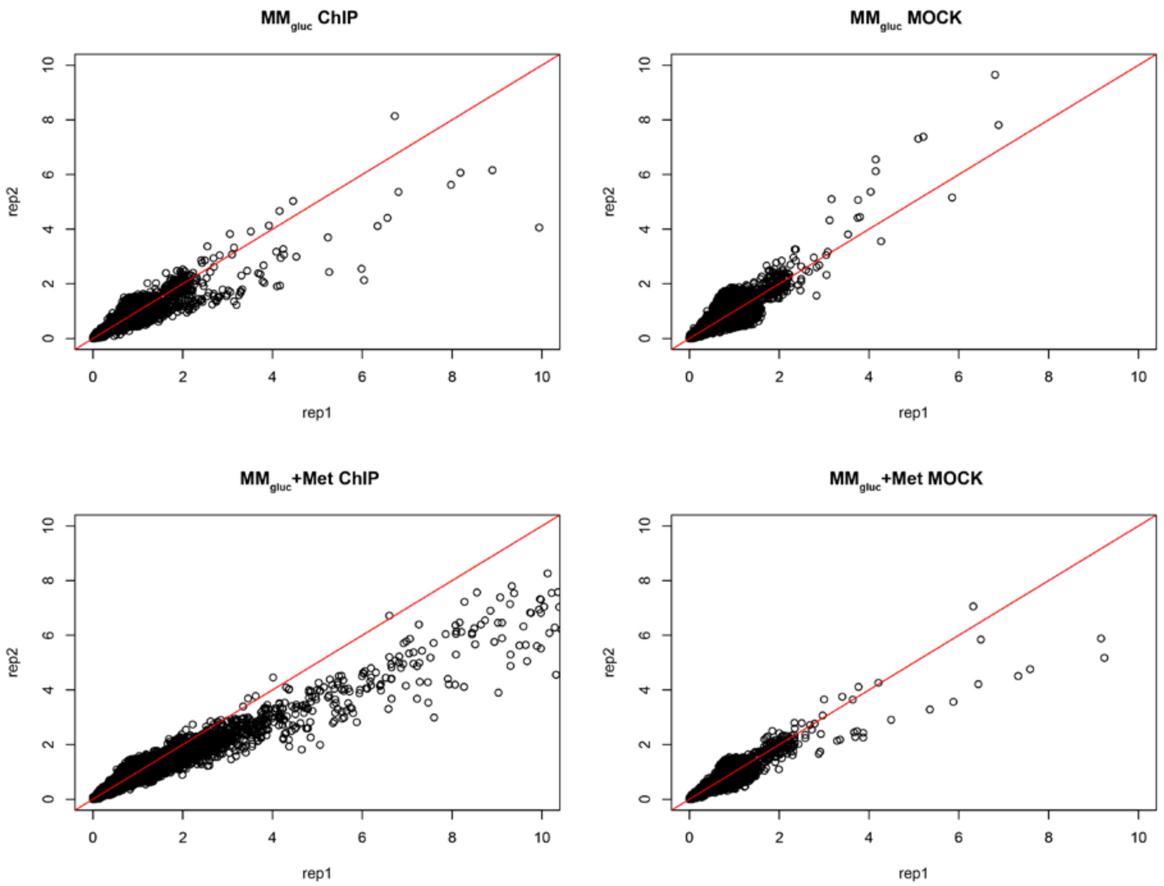


**Figure S2: Gcn4 protein expression, and ChIP-seq correlation plots**
A. A representative western blot for Gcn4-HA (Gcn4 tagged with the HA epitope at the endogenous locus) expression in the indicated medium. Gcn4 protein levels are high in $MM_{gluc}$+Met. Abbreviations: RM - rich medium, $MM_{gluc}$ - minimal medium without amino acids and with glucose as a carbon source, and $MM_{gluc}$+Met - minimal medium without amino acids and with glucose as a carbon source, supplemented with 2mM methionine.
B. Correlation plots between normalised read counts of the ChIP sequencing data, from cells growin in different medium. Normalised read counts of 100000 randomly selected positions from the genome are plotted. ChIP data show clear enrichment/DNA binding for Gcn4 only in methionine supplemented conditions. Replicates shows good correlation with a Pearson correlation coefficient, R = 0.9.

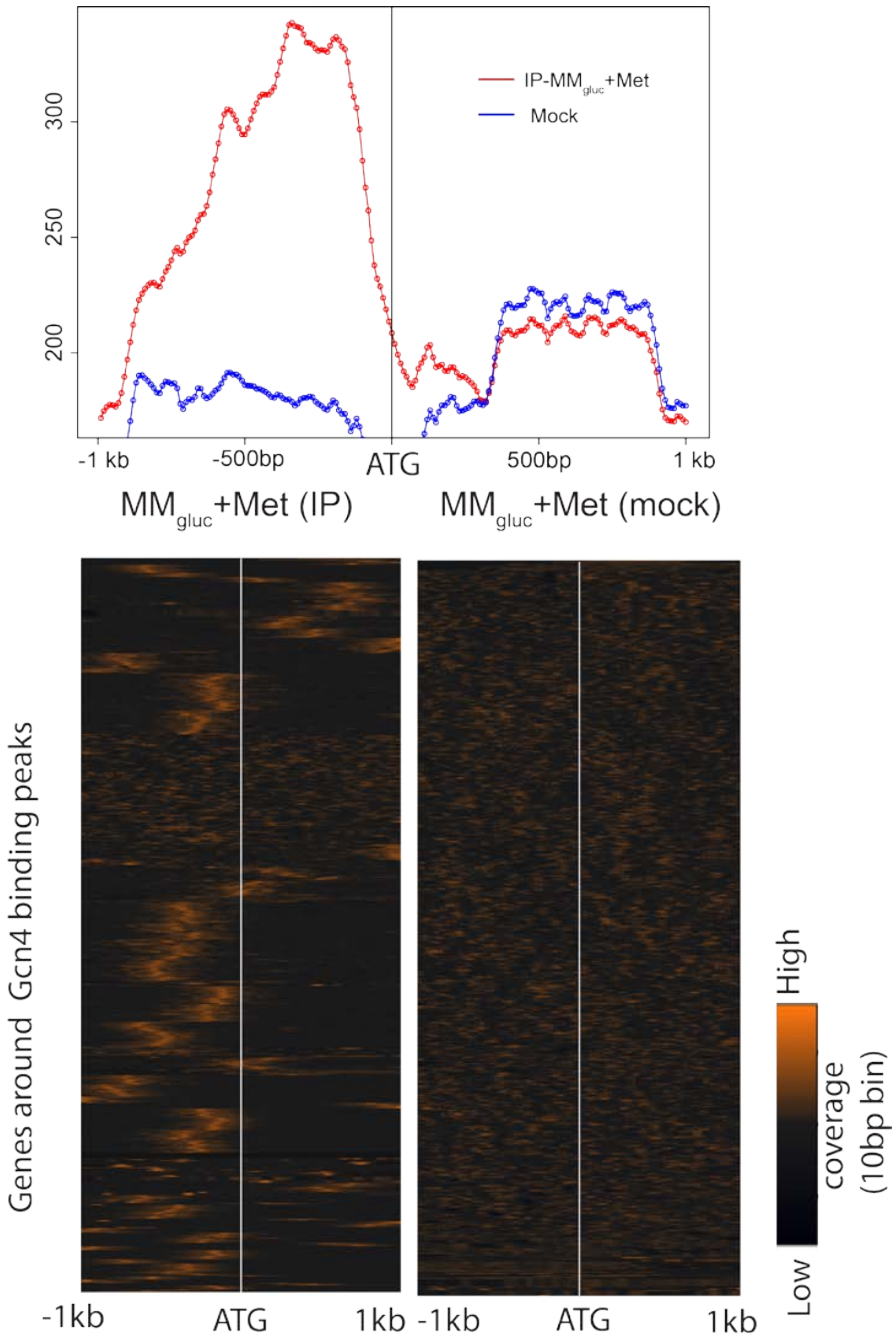


**Figure S3: Read coverage for Gcn4 binding, in the context of predicted translation sites of transcribed mRNAs.**

A heatmap showing the read coverage from 1kb bases flanking the predicted translation start site (ATG) of genes, for the Gcn4 targets. The heatmap on the left shows read coverage in ChIP samples, and on the right shows coverage in mock-ChIP (control) in the $MM_{gluc}$ +Met condition. The density plot in the top panel shows that most of the target genes have Gcn4 binding peaks upstream of the translation start site in ChIP samples (red) and such a pattern is absent in mock samples (blue). The deepTools package was used for generating the matrix in this analysis (Ramírez et al., 2016).

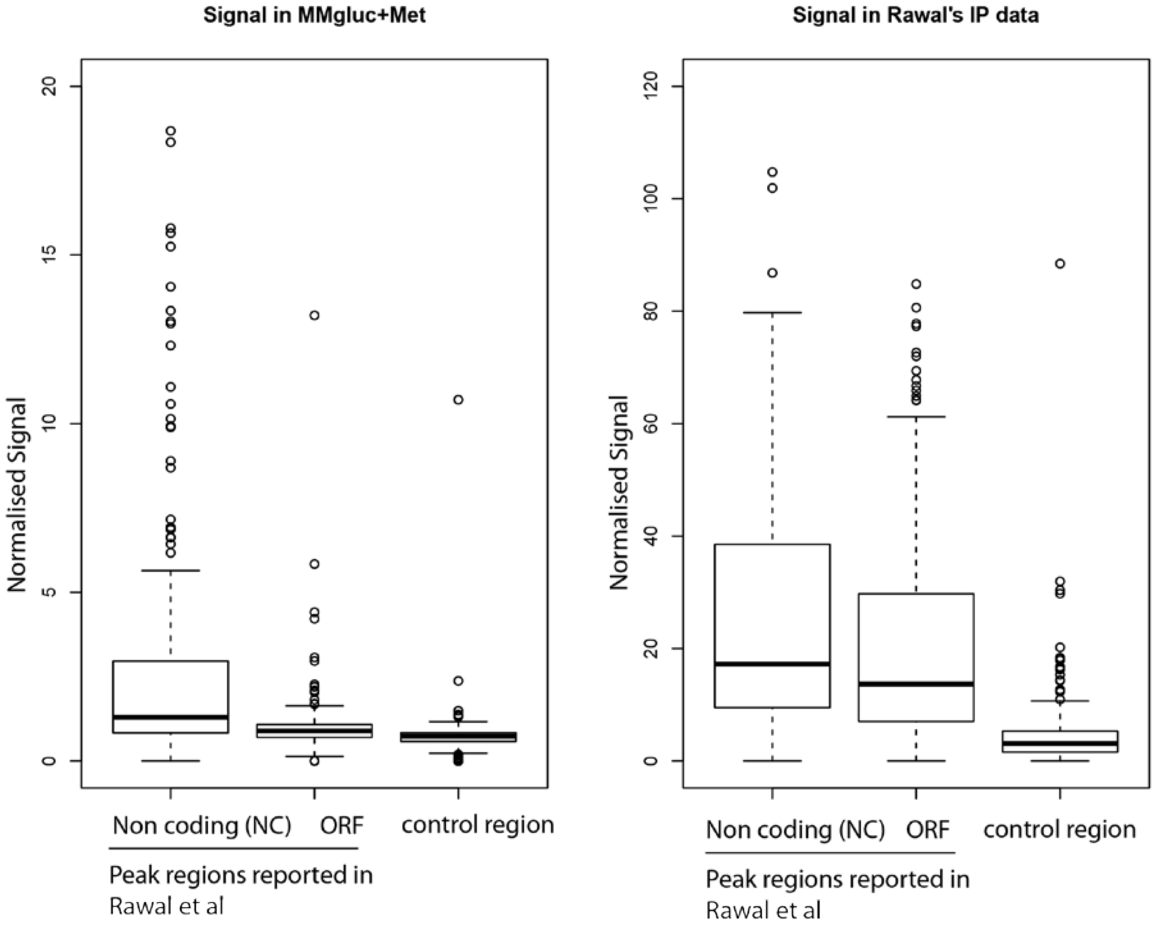


**Figure S4: Gcn4 binding signal is high in non-coding regions relative to ORF regions, in $MM_{gluc}$+Met medium:**

For Gcn4 binding in the non-coding and the open reading frame (ORF) regions, as reported in a previous study (Rawal et al., 2018) we calculated the Gcn4 binding signal in the Gcn4 ChIP sequencing data from the $MM_{gluc}$+Met condition (current study). Notably, in the $MM_{gluc}$+Met condition, the Gcn4 binding signal in ORF peaks is significantly lower than the non-coding region peaks ($p<10e-8$). Contrarily, in the previous study (Rawal et al., 2018), the Gcn4 binding signal found in ORF and non-coding regions are very similar. Therefore, we confirm that the most of the peaks found in our Gcn4 ChIP, in the $MM_{gluc}$+Met, are present in non-coding regions of the genome, and not in the ORFs of genes.

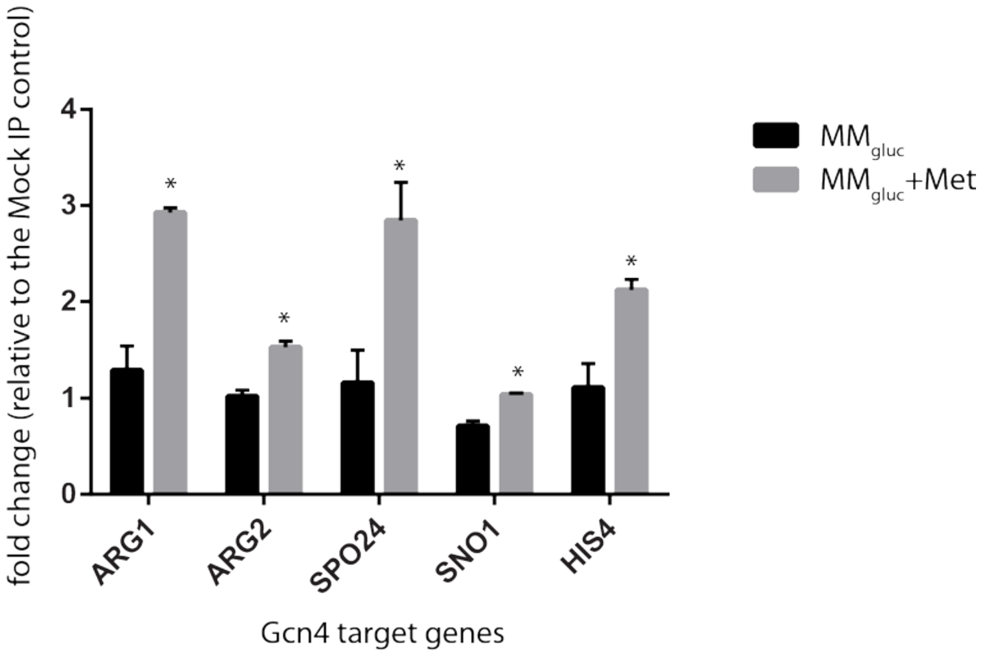


**Figure S5**

ChIP qPCR validation of selected Gcn4 binding targets. Gcn4 binding targets identified in the $MM_{gluc}$+Met condition were further confirmed using a traditional quantitative PCR, with the primers specific to the promoter region of the target genes. Gcn4 binding to the promoter of these targets is significantly high in $MM_{gluc}$+Met, relative to the $MM_{gluc}$. Data shown are mean±SD from 2 biological replicates. * p-value < 0.05.

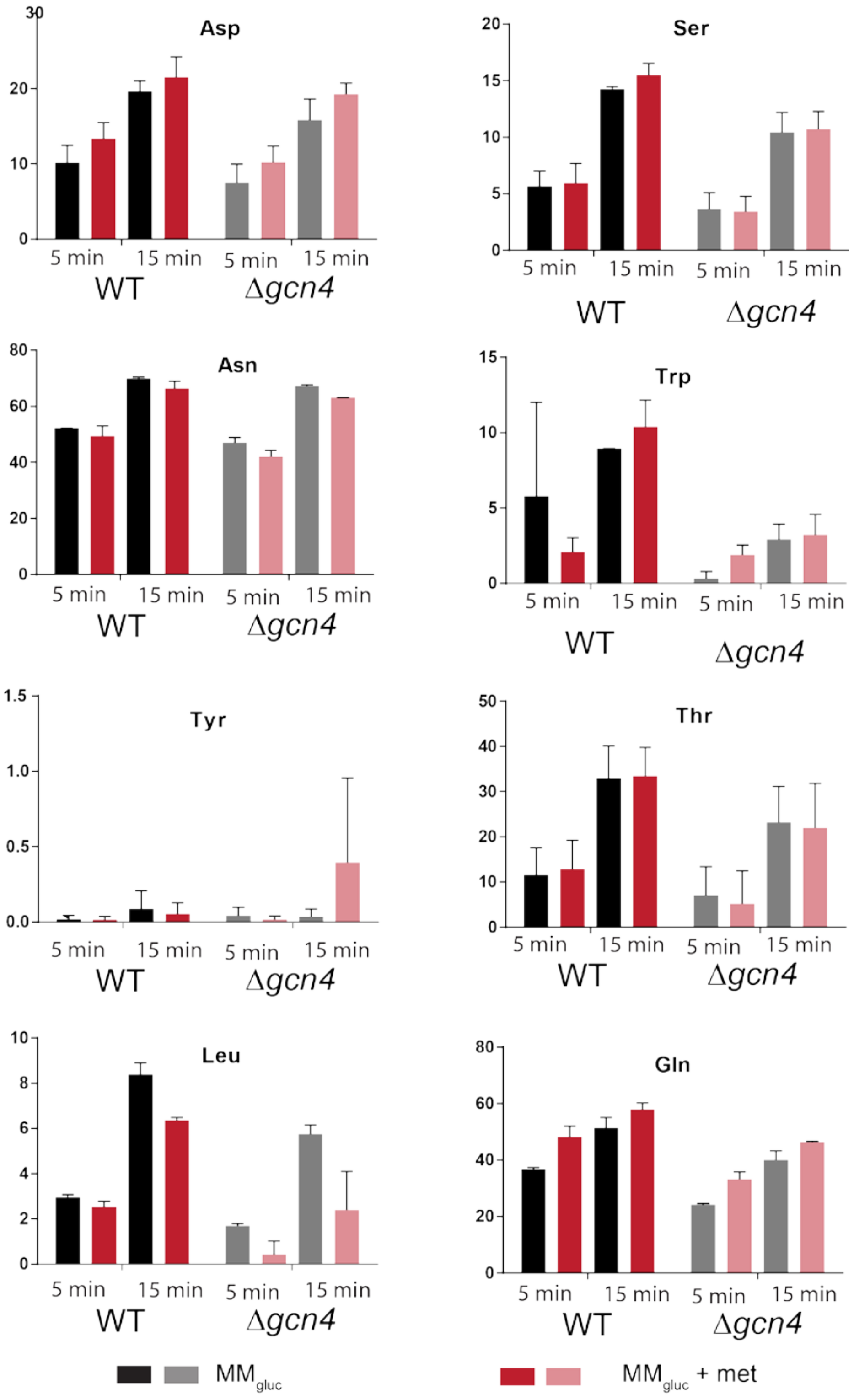


**Figure S6 Quantitative LC/MS/MS based metabolic flux analysis, using $N^{15}$ ammonium sulfate labelling to estimate new amino acid synthesis in a methionine and Gcn4 dependent manner.**

WT and *Δgcn4* cells grown in RM were shifted to $MM_{gluc}$ or $MM_{gluc}$+Met with 0.5X of ammonium sulfate, and 0.5X of $N^{15}$ labelled ammonium sulfate, and metabolites extracted from cells 5 and 15 minutes after the pulse. Label incorporation into the indicated amino acids was measured using LC-MS/MS analysis.

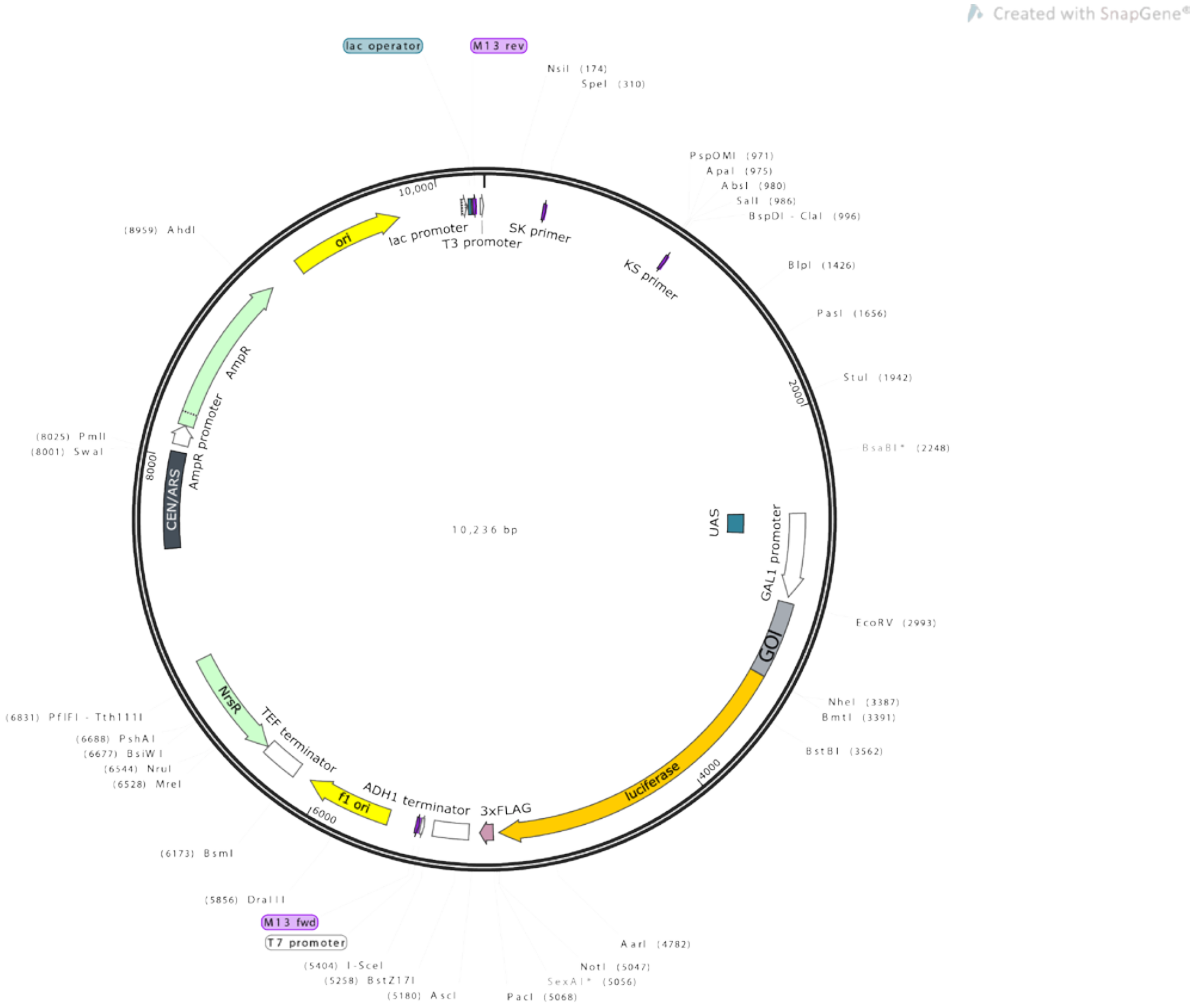


**Figure S7 Inducible luciferase-conjugated reporters to estimate expression of lysine and arginine enriched genes.**

The complete open reading frames (ORFs) of the indicated genes were cloned in place of 'GOI-Gene of Interest' labelled in the plasmid map. In this study, ORFs of genes *RPL32*, *NHP2*, *STM1*, *NHP2*, *RPS20* were cloned upstream of the luciferase sequence, and the resulting plasmids were named pSL218, pSL221, pSL224, pSL234 respectively. In this system, the expression of the GOI+luciferase is under the control of an inducible promoter, which is induced upon adding ß- estradiol, leading to gene expression and translation. The activity of luciferase enzyme (which is cloned in frame with the gene of interest (GOI)), serves as a quantitative indicator of the translation of that reporter in the given condition and genetic background. The plasmid map was created using SnapGene viewer version 3.3.4 (http://www.snapgene.com)

**Supplementary table 1**

| Strains | Genotype | Reference |
|---|---|---|
| WT (CEN.PK) | Mat a | (J.P. van Dijken et al., 2000) |
| Gcn4-HA | MAT a, *GCN4-HA-KanMX* | (Walvekar et al., 2018) |
| Gcn4-FLAG | MAT a, *GCN4-FLAG-KanMX* | This Study |
| *gcn4Δ::NAT* | MAT a, *gcn4Δ::NAT* | (Walvekar et al., 2018) |
| *gcn4Δ::KanMX* | MAT a, *gcn4Δ::KanMX* | This Study |

**Supplementary table 2**

| Plasmid | Promoter and Reporter | Reference |
|---|---|---|
| pGL3-basic | | Addgene |
| pGEV-FLAG | GEV- estradiol inducible promoter | (McIsaac et al., 2011) |
| pSL207 (modified version of pGEV-FLAG) | GEV- estradiol inducible promoter | This study |
| pSL217 | *pGEV-Luciferase::NAT* | This study |
| pSL218 | *pGEV-RPL32-Luciferase::NAT* | This study |
| pSL221 | *pGEV-NHP2-Luciferase::NAT* | This study |
| pSL224 | *pGEV-STM1-Luciferase::NAT* | This study |
| pSL234 | *pGEV-RPS20-Luciferase::NAT* | This study |

**Supplementary References**

J.P. van Dijken, J.B., Brambillac, L., Dubocd, P., Francoise, J.M., Gancedof, C., Giusepping, M.L.F.HEijnenh, J.J., Hoarei, M., Langej, H.C., Maddenk, E.A., Niederbergerb, P., et al. (2000). An interlaboratory comparison of physiological and genetic properties of four Saccharomyces cerevisiae strains. Enzyme Microb. Technol. *26*, 706–714.

McIsaac, R.S., Silverman, S.J., McClean, M.N., Gibney, P.A., Macinskas, J., Hickman, M.J., Petti, A.A., and Botstein, D. (2011). Fast-acting and nearly gratuitous induction of gene expression and protein depletion in Saccharomyces cerevisiae. Mol. Biol. Cell *22*, 4447–4459.

Ramírez, F., Ryan, D.P., Grüning, B., Bhardwaj, V., Kilpert, F., Richter, A.S., Heyne, S., Dündar, F., and Manke, T. (2016). deepTools2: a next generation web server for deep-sequencing data analysis. Nucleic Acids Res. *44*, W160–W165.

Rawal, Y., Chereji, R. V, Valabhoju, V., Qiu, H., Ocampo, J., Clark, D.J., and Hinnebusch, A.G. (2018). Gcn4 Binding in Coding Regions Can Activate Internal and Canonical 5' Promoters in Yeast. Mol. Cell *70*, 297-311.e4.

Walvekar, A.S., Srinivasan, R., Gupta, R., and Laxman, S. (2018). Methionine coordinates a hierarchically organized anabolic program enabling proliferation. Mol. Biol. Cell *29*, 3183–3200.
